# Detection of Alzheimer’s associated microRNAs using a DNA-based smart reagent

**DOI:** 10.1101/2021.06.01.446618

**Authors:** Arun Richard Chandrasekaran, Ken Halvorsen

## Abstract

Alzheimer’s disease (AD) is the most common neurodegenerative disorder, with significant research efforts devoted to identifying new biomarkers for clinical diagnosis and treatment. MicroRNAs have emerged as likely disease regulators and biomarkers for AD, now implicated as having roles in several biological processes related to progression of the disease. In this work, we use the *miRacles* assay (**mi**cro**R**NA **a**ctivated **c**onditional **l**ooping of **e**ngineered **s**witches) for single-step detection of AD-related microRNAs. The technology is based on conformationally responsive DNA nanoswitches that loop upon recognition of a target microRNA and report their on/off status through an electrophoretic readout. Unlike many other methods, our approach directly detects native microRNAs without amplification or labeling, eliminating the need for expensive enzymes, reagents, and equipment. We used this assay to screen for AD-related microRNAs, demonstrate specificity within a microRNA family, sensitivity of ∼ 8 fM, and multiplexing capability to simultaneously detect four microRNA targets. Toward clinical use, we provide proof-of-concept detection and quantifiable dysregulation of specific microRNAs from total RNA extracts derived from healthy and AD brain samples. In the context of AD, this “smart reagent” could facilitate biomarker discovery, accelerate efforts to understand the role of microRNAs in AD, and have clinical potential as a diagnostic or monitoring tool for validated biomarkers.

---

Alzheimer’s disease (AD) is a neurodegenerative disorder characterized by memory loss and multiple cognitive abnormalities.^1^ With no cure and minimal early detection, there are significant research efforts into all aspects of the disease spanning from basic disease biology to clinical diagnosis and treatment. Typical diagnosis for AD relies on cognitive assessment,^2^ brain magnetic resonance imaging (MRI),^3^ and detection of cerebrospinal fluid tau protein and amyloid biomarkers.^4^ There have also been recent efforts at creating deep learning models using a combination of imaging, genetic analysis, and clinical test data from such diagnostic methods.^5^ More recently, several approaches have been developed for highly sensitive detection of Aβ and tau in the blood.^6^ However, these biomarkers alone may not reflect the complex molecular pathophysiology of AD, creating a need for studying changes in other biomarkers such as synaptic markers, metabolites and microRNAs.^7^

MicroRNAs (short, noncoding, regulatory RNAs) are implicated to have roles in several biological processes related to progression of AD.^8–10^ The differential levels of certain microRNAs in circulating fluids (blood, serum, plasma, exosomes, cerebrospinal fluid) could be used as potential biomarkers for early detection of AD.^11,12^ Beyond the conventional microRNA detection methods (qRT-PCR, microarrays, Next Generation Sequencing and Northern blotting), several new methods based on nanomaterials such as graphene, carbon nanotubes, metallic nanoparticles and quantum dots have been developed.^13–15^ DNA nanotechnology has also contributed to this development, with structures such as DNA tetrahedra, DNA origami scissors, plasmonic DNA-gold nanorods and DNA walkers used for microRNA detection.^16^ In some of these methods, the structures are overly complex to construct, need multiple steps, enzyme-based amplification, expensive equipments and time-consuming strategies for readout. More development is needed to obtain direct, unamplified microRNA detection with easily adaptable readout methods.

Working towards this goal, we present a low cost, non-enzymatic, direct microRNA detection assay based on DNA nanoswitches for the detection of AD-related microRNAs termed *miRacles* (**mi**cro**R**NA **a**ctivated **c**onditional **l**ooping of **e**ngineered **s**witches).^17^ We previously used the DNA nanoswitches for detecting nucleic acids (DNA,^18^ microRNAs,^17^ viral RNAs^19^) and other biomarkers such as proteins^20^ and enzymes.^21^ In this work, we use the nanoswitches to detect AD-related microRNAs with femtomolar sensitivity, specificity to discriminate microRNAs from the same family and demonstrate multiplexed detection of four microRNAs in a single assay. We further screened for AD-related microRNAs in brain tissue total RNA from healthy and AD samples, and quantified dysregulation of specific microRNA expression levels in these samples.

The DNA nanoswitch is constructed based on the principles of DNA origami,^22^ and designed to be a long duplex comprised of a single-stranded scaffold (commercially available M13 viral genome) and short complementary backbone oligonucleotides (**Figure 1a** and **Figure S1**). Two of the backbone oligonucleotides are modified to contain single stranded extensions that act as detectors to capture a target microRNA. On binding the microRNA, the nanoswitch is reconfigured from its linear (off) state to a looped (on) state, and the two states can be easily identified on an agarose gel (**Figure 1b**). The presence of the “on” band indicates the presence of the target microRNA in the sample. The highlight of this assay is that it does not require any additional labeling or amplification; the signal comes from the nanoswitch itself, through the intercalation of thousands of dye molecules on the nanoswitch (using regular DNA gel stains such as GelRed).

**Figure 1.**
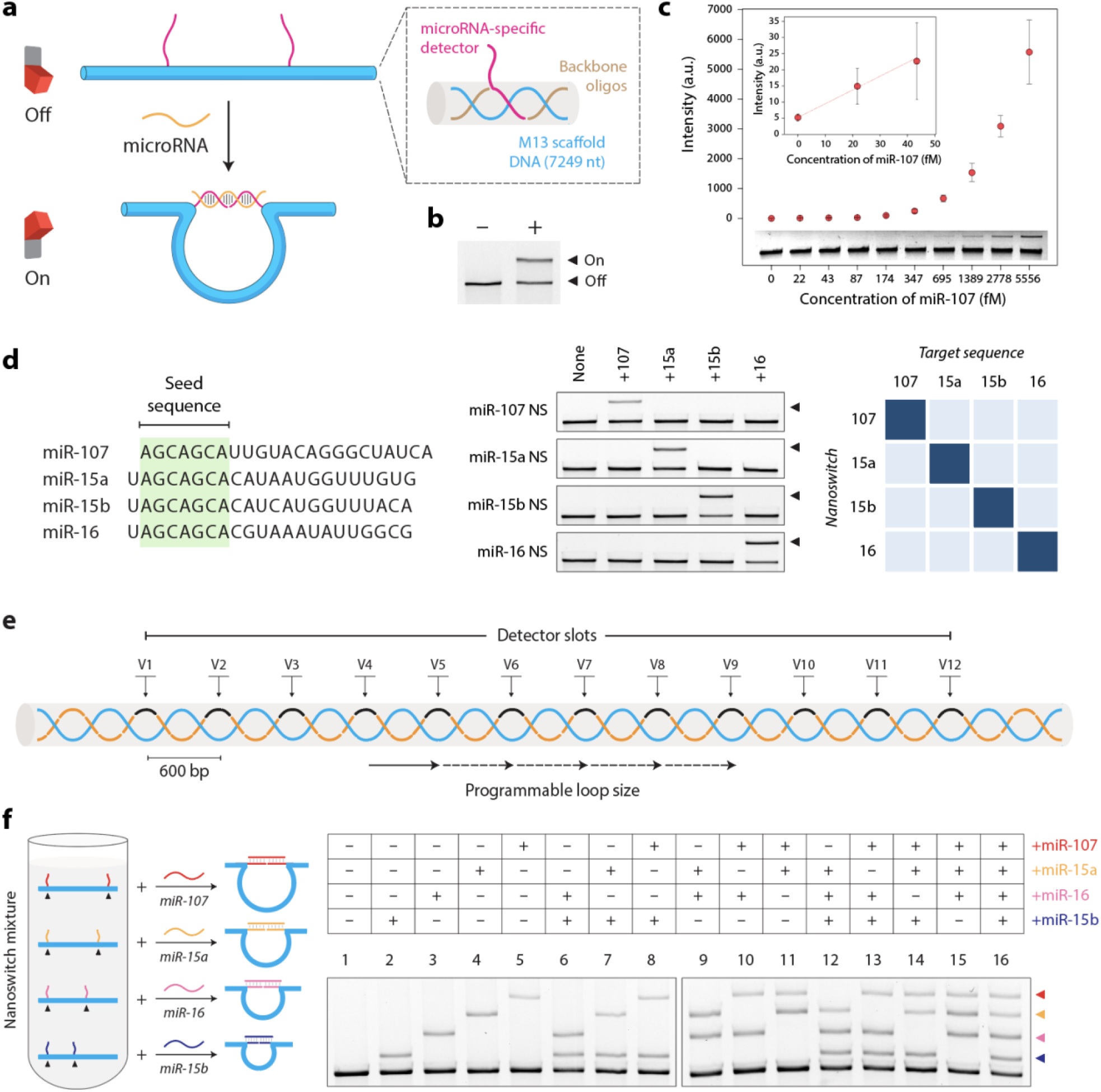
Design and characterization of nanoswitch assay for Alzheimer’s disease microRNAs. (a) The DNA nanoswitch is a linear duplex constructed from single stranded M13 scaffold DNA and short complementary backbone oligonucleotides (inset). Two single stranded extensions act as detectors to bind complementary portions on a microRNA target. The binding event reconfigures the nanoswitch from the linear “off” state to the looped “on” state. (b) The two states can be easily identified on an agarose gel for an easy readout of microRNA detection. (c) Sensitivity plot and corresponding gel image (inset) for miR-107. Data points and error bars represent the means and SD, respectively, from triplicate measurements. (d) Specificity of the assay for the miR-107 super family of microRNAs which share the same seed sequence. Specificity of each nanoswitch to its target is shown by the gel image and heat map corresponding to normalized band intensities. (e) Variable regions (V1−V12) on the DNA nanoswitch allow programmable placement of detectors, resulting in nanoswitches with different loop sizes. (f) Four-channel multiplexing using a nanoswitch mixture containing nanoswitches of different loop sizes. Each nanoswitch provides a unique signal for each microRNA target present. Gel image shows detection of all 16 combinations of four AD-related microRNAs.

For validating AD-related microRNA detection, we chose miR-107 as an example, which is known to accelerate AD progression by multiply targeting the 3′-untranslated region (UTR) of β-site amyloid precursor protein-cleaving enzyme 1 (BACE1) mRNA.^23^ We performed sensitivity experiments by analyzing nanoswitch looping with different amounts of miR-107 and showed visible detection to 175 fM and a calculated limit of detection (LOD) of ∼7.7 fM (defined as the concentration of biomarker that yields a signal exceeding the mean background by 3 SDs) (**Figure 1c** and **Figure S2**). This sensitivity is higher than some of the highly expressed microRNAs reported in brain samples.^24^

Next, we investigated the ability of our nanoswitches to distinguish closely-related sequences. This is important as naturally occurring microRNAs can sometimes contain similar sequences, with such microRNAs being classified into the same family. As an example, we chose the miR-107 superfamily of microRNAs in which miR-107, miR-15a/b and miR-16 have been documented to be dysregulated in AD brain.^25^ mir-16 is also known to be a post-translational regulator of amyloid protein precursor, and its low expression leads to amyloid-beta (Aβ) deposition, leading to high risk of AD.^26^ These microRNAs share the same seed sequence (**Figure 1d**) and thus detection assays need to be able to distinguish between these closely-related microRNAs. We designed nanoswitches with detectors specific to microRNAs 107, 15a, 15b and 16, and tested specificity of detecting these microRNAs. Results showed perfect specificity, with each nanoswitch detecting only its corresponding target and not the others (gels and heat map in **Figure 1d**). Along with our previously reported single-nucleotide specificity,^17^ these results illustrate the high specificity of our assay which has been a key challenge for microRNA detection.^27^

For diagnostics, detection of biomarker panels has been shown in some cases to be more accurate than individual biomarkers.^28^ We used the programmability of the nanoswitches to develop a multiplexing system capable of detecting multiple microRNAs from the same sample. We positioned detector strands on specific locations on the DNA scaffold, resulting in loops of different sizes when bound to the target RNA (**Figure 1e**). The loop size of the nanoswitch determines the gel migration, with each loop size reported as a unique band on the gel. To demonstrate multiplexing, we chose the same four microRNAs we used for specificity (microRNAs 107, 15a, 15b and 16) and designed nanoswitches for each microRNA with different loop sizes (**Figure 1f**). For multiplexed assay, we created a nanoswitch mix containing these four nanoswitches and detected each microRNA (**Figure 1f**, lanes 2-5), subset of two (lanes 6-11), subset of three (lanes 12-15) and all four targets in a single gel lane (lane 16). This multiplexing strategy enables direct comparison of microRNA levels in one sample without labeling or amplification, and also moves a step towards expanding the throughput of the *miRacles* assay.

To demonstrate biological relevance, we then screened for AD-related microRNAs in total RNA samples derived from healthy and AD brain tissues (**Figure 2a**). We selected a set of AD-related microRNAs reported in the literature^10,23,29–34^ and tested corresponding DNA nanoswitches for detection of these microRNAs in 500 ng of healthy and AD brain total RNA extracts (**Figure S3**). From the obtained “hits”, we analyzed miR-125b, miR-100-5p, miR-29a-3p and miR-29b-3p as representative examples, showing different levels of upregulation or downregulation when comparing healthy and AD samples (**Figure S4**). Specifically, miR-29a and miR-29b are known to be downregulated as disease progresses in AD patients.^35,36^ We chose these two microRNAs to quantitatively evaluate differential expression in healthy and AD samples using total RNA from different brain tissues such as frontal, parietal and temporal lobes along with full brain total RNA extracts (**Figure 2b**). Before testing in total RNA, we ensured that there is no crosstalk in detecting miR-29a-3p and miR-29b-3p using DNA nanoswitches since they have similar sequences (**Figure S5**). For detecting from brain total RNA, we incubated DNA nanoswitches with 500 ng of total RNA extracts from different brain tissues and observed different levels of detection in each case for miR-29a-3p and miR-29b-3p (**Figure 2c, 2f and Figure S6-S7**). We quantified the signal intensities and found different levels of dysregulation of both these microRNAs in the AD samples (**Figure 2d, 2g**). We also calculated the fold change in expression of these microRNAs and found that both miR-29a-3p and miR-29b-3p were downregulated in total RNA samples from all the AD brain tissue types (**Figure 2e, 2h**).

**Figure 2.**
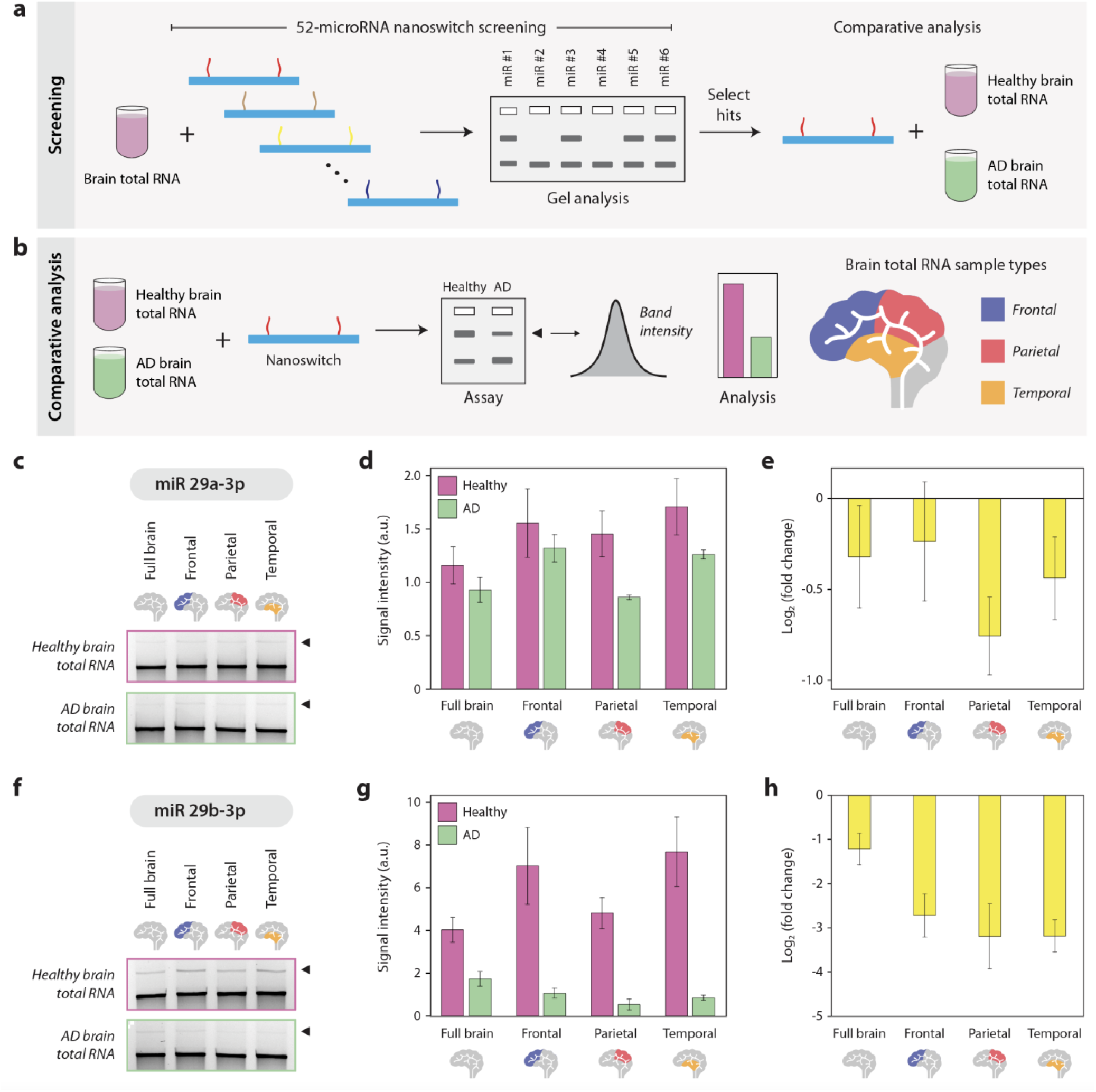
Detection and quantitation of microRNAs from brain total RNA samples. (a) Workflow of AD-related microRNA screening using DNA nanoswitches. (b) Schematic of comparative analysis of dysregulated microRNAs in healthy and AD brain total RNA samples. (c) Gel images showing detection of miR-29a-3p in total RNA extracts of healthy and AD brain tissues. (d) Quantified signal intensity of miR-29a-3p from gels in (b) showing downregulation in AD samples. (e) Fold change in expression of miR-29a-3p in AD brain total RNA compared to healthy samples. (f, g, h) Gel images, detection signal and fold change in expression of miR-29b-3p in healthy and AD brain total RNA. Data points and error bars represent the means and SD, respectively, from triplicate measurements.

Given the ability to detect and quantify changes in individual microRNAs, we then created a multiplexed assay to detect both miR-29a-3p and miR-29b-3p in one assay from a single brain total RNA sample. We designed nanoswitches for miR-29a-3p and miR-29b-3p with different loop sizes and created a nanoswitch mix (**Figure 3a**). Using this mix, we detected both miR-29a-3p and miR-29b-3p from a single healthy or AD brain total RNA sample (**Figure 3a, inset and Figure S8**). Quantified results again showed downregulation of both microRNAs (**Figure 3b**), with similar fold changes in expression compared to when the microRNAs were assayed individually (**Figure 3c**).

**Figure 3.**
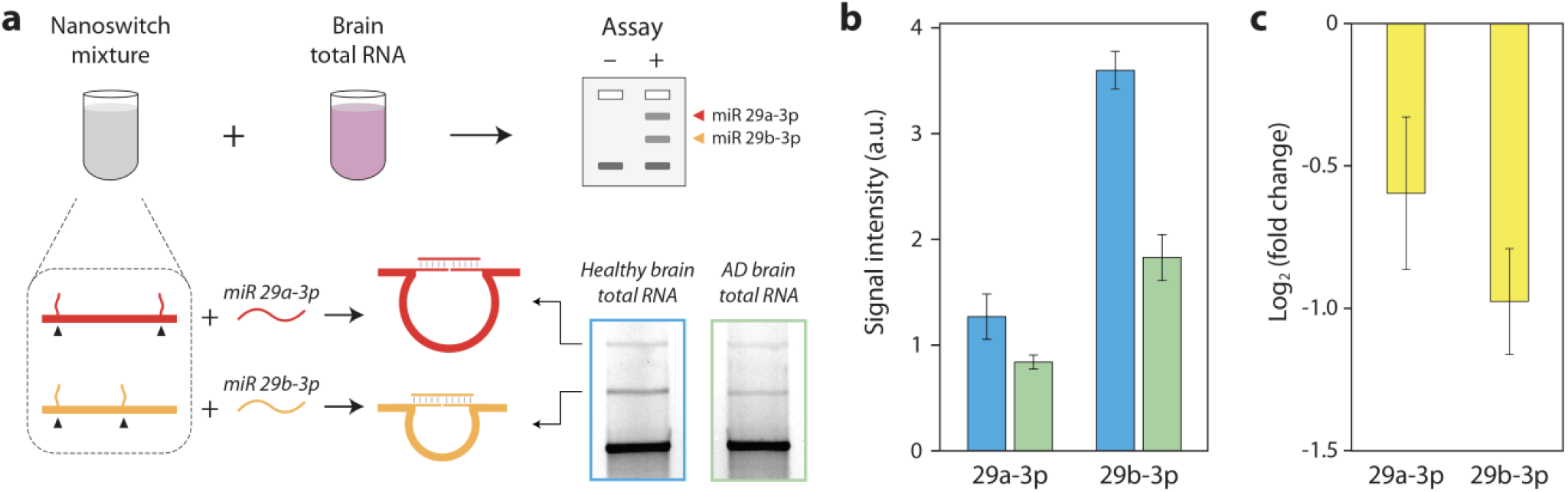
Multiplexed analysis in brain total RNA. (a) A nanoswitch mix consisting of two nanoswitches with different loop sizes, each corresponding to miR-29a-3p or miR-29b-3p can be used to detect both microRNAs in a single assay. Inset shows gel images of microRNA detection from healthy and AD brain samples. (b) Quantified signal intensity of miR-29a-3p and miR-29b-3p from gels shown in (a). (c) Fold change in expression of miR-29a-3p and miR-29b-3p in AD brain total RNA compared to healthy samples. Data points and error bars represent the means and SD, respectively, from triplicate measurements.

In this study, we show that a DNA nanoswitch can be used as a “smart reagent” for sensitive, specific and quantifiable detection of AD-related microRNAs. Our programmable nanoswitch reports molecular recognition events by transforming them into topological transitions that can be easily read out using the common lab technique of gel electrophoresis. Recognizing that barriers of cost and complexity slow new technology adoption, our DNA nanoswitches repurpose an existing and familiar lab tool, maximizing accessibility. Compared to existing techniques of nucleic acid detection, our *miRacles* assay is low-cost (<10 ¢/lane, **Table S1**), can be combined with portable e-gels, and does not require amplification, enzymes or expensive equipment. Further, the nanoswitches can be dried and stored for use^37^ enabling potential point-of-care use. The programmability of DNA nanoswitches provides flexibility in design, especially in multiplexing, enabling placement of multiple detectors in the same sensor that can be independently resolved. This enables comparative analysis from one sample, which is becoming increasingly important as evidence points to biomarker panels as being more effective than individual biomarkers for clinical applications,^28^ and researchers struggle with problems of data normalization in biosensing.^38^ Further, analysis of microRNA panels may report multiple pathways contributing to AD pathology, enabling the design of personalized therapies.^32^ Increasing evidence that plasma or serum-based biomarkers can potentially distinguish different neurodegenerative conditions such as AD, Parkinson’s disease and schizophrenia makes our assay potentially useful as a diagnostic tool for neurological diseases.^39^ Our approach can be applied to AD-related microRNAs in research as an alternative to qPCR and Northern blotting, with faster microRNA screening methods being useful in drug discovery.^25^ Ultimately, our approach also has potential to identify and measure circulating or exosomal microRNAs^40^ as AD biomarkers for early detection as well as monitoring at-risk patients.

## Acknowledgments

Research reported in this publication was supported by the NIH through NIGMS and NIA under award R35GM124720 and R35GM124720-03S1 to K.H. The authors thank Andrew Hayden and Javier Vilcapoma for technical support.

## Notes

A. R. C. and K. H. are inventors on patents and patent applications covering aspects of the DNA nanoswitch design and applications.

## Author contributions

A.R.C designed and performed experiments, analyzed and visualized data and wrote the first draft of the manuscript. K.H. conceived the project, designed experiments and co-wrote later drafts of the manuscript.

## Reference

(1) Mucke, L. Alzheimer’s Disease. Nature 2009, 461 (7266), 895–897. https://doi.org/10.1038/461895a.

(2) Loewenstein, D. A.; Curiel, R. E.; Duara, R.; Buschke, H. Novel Cognitive Paradigms for the Detection of Memory Impairment in Preclinical Alzheimer’s Disease. Assessment 2018, 25 (3), 348–359. https://doi.org/10.1177/1073191117691608.

(3) Acharya, U. R.; Fernandes, S. L.; WeiKoh, J. E.; Ciaccio, E. J.; Fabell, M. K. M.; Tanik, U. J.; Rajinikanth, V.; Yeong, C. H. Automated Detection of Alzheimer’s Disease Using Brain MRI Images– A Study with Various Feature Extraction Techniques. J. Med. Syst. 2019, 43 (9), 302. https://doi.org/10.1007/s10916-019-1428-9.

(4) Blennow, K.; Mattsson, N.; Schöll, M.; Hansson, O.; Zetterberg, H. Amyloid Biomarkers in Alzheimer’s Disease. Trends Pharmacol. Sci. 2015, 36 (5), 297–309. https://doi.org/10.1016/j.tips.2015.03.002.

(5) Venugopalan, J.; Tong, L.; Hassanzadeh, H. R.; Wang, M. D. Multimodal Deep Learning Models for Early Detection of Alzheimer’s Disease Stage. Sci. Rep. 2021, 11 (1), 3254. https://doi.org/10.1038/s41598-020-74399-w.

(6) Nabers, A.; Perna, L.; Lange, J.; Mons, U.; Schartner, J.; Güldenhaupt, J.; Saum, K.-U.; Janelidze, S.; Holleczek, B.; Rujescu, D.; Hansson, O.; Gerwert, K.; Brenner, H. Amyloid Blood Biomarker Detects Alzheimer’s Disease. EMBO Mol. Med. 2018, 10 (5), e8763. https://doi.org/10.15252/emmm.201708763.

(7) Khoury, R.; Ghossoub, E. Diagnostic Biomarkers of Alzheimer’s Disease: A State-of-the-Art Review. Biomark. Neuropsychiatry 2019, 1, 100005. https://doi.org/10.1016/j.bionps.2019.100005.

(8) Delay, C.; Mandemakers, W.; Hébert, S. S. MicroRNAs in Alzheimer’s Disease. Neurobiol. Dis. 2012, 46 (2), 285–290. https://doi.org/10.1016/j.nbd.2012.01.003.

(9) Reddy, P. H.; Tonk, S.; Kumar, S.; Vijayan, M.; Kandimalla, R.; Kuruva, C. S.; Reddy, A. P. A Critical Evaluation of Neuroprotective and Neurodegenerative MicroRNAs in Alzheimer’s Disease. Biochem. Biophys. Res. Commun. 2017, 483 (4), 1156–1165. https://doi.org/10.1016/j.bbrc.2016.08.067.

(10) Hu, Y.-B.; Li, C.-B.; Song, N.; Zou, Y.; Chen, S.-D.; Ren, R.-J.; Wang, G. Diagnostic Value of MicroRNA for Alzheimer’s Disease: A Systematic Review and Meta-Analysis. Front. Aging Neurosci. 2016, 8. https://doi.org/10.3389/fnagi.2016.00013.

(11) Kumar, P.; Dezso, Z.; MacKenzie, C.; Oestreicher, J.; Agoulnik, S.; Byrne, M.; Bernier, F.; Yanagimachi, M.; Aoshima, K.; Oda, Y. Circulating MiRNA Biomarkers for Alzheimer’s Disease. PLOS ONE 2013, 8 (7), e69807. https://doi.org/10.1371/journal.pone.0069807.

(12) Kiko, T.; Nakagawa, K.; Tsuduki, T.; Furukawa, K.; Arai, H.; Miyazawa, T. MicroRNAs in Plasma and Cerebrospinal Fluid as Potential Markers for Alzheimer’s Disease. J. Alzheimers Dis. 2014, 39 (2), 253–259. https://doi.org/10.3233/JAD-130932.

(13) Graybill, R. M.; Bailey, R. C. Emerging Biosensing Approaches for MicroRNA Analysis. Anal. Chem. 2016, 88 (1), 431–450. https://doi.org/10.1021/acs.analchem.5b04679.

(14) Pritchard, C. C.; Cheng, H. H.; Tewari, M. MicroRNA Profiling: Approaches and Considerations. Nat. Rev. Genet. 2012, 13 (5), 358–369. https://doi.org/10.1038/nrg3198.

(15) Dave, V. P.; Ngo, T. A.; Pernestig, A.-K.; Tilevik, D.; Kant, K.; Nguyen, T.; Wolff, A.; Bang, D. D. MicroRNA Amplification and Detection Technologies: Opportunities and Challenges for Point of Care Diagnostics. Lab. Invest. 2019, 99 (4), 452–469. https://doi.org/10.1038/s41374-018-0143-3.

(16) Chandrasekaran, A. R.; Punnoose, J. A.; Zhou, L.; Dey, P.; Dey, B. K.; Halvorsen, K. DNA Nanotechnology Approaches for MicroRNA Detection and Diagnosis. Nucleic Acids Res. 2019, 47 (20), 10489–10505. https://doi.org/10.1093/nar/gkz580.

(17) Chandrasekaran, A. R.; MacIsaac, M.; Dey, P.; Levchenko, O.; Zhou, L.; Andres, M.; Dey, B. K.; Halvorsen, K. Cellular MicroRNA Detection with MiRacles: MicroRNA-Activated Conditional Looping of Engineered Switches. Sci. Adv. 2019, 5 (3), eaau9443. https://doi.org/10.1126/sciadv.aau9443.

(18) Chandrasekaran, A. R.; Zavala, J.; Halvorsen, K. Programmable DNA Nanoswitches for Detection of Nucleic Acid Sequences. ACS Sens. 2016, 1 (2), 120–123. https://doi.org/10.1021/acssensors.5b00178.

(19) Zhou, L.; Chandrasekaran, A. R.; Punnoose, J. A.; Bonenfant, G.; Charles, S.; Levchenko, O.; Badu, P.; Cavaliere, C.; Pager, C. T.; Halvorsen, K. Programmable Low-Cost DNA-Based Platform for Viral RNA Detection. Sci. Adv. 2020, eabc6246. https://doi.org/10.1126/sciadv.abc6246.

(20) Chandrasekaran, A. R.; MacIsaac, M.; Vilcapoma, J.; Hansen, C. H.; Yang, D.; Wong, W. P.; Halvorsen, K. DNA Nanoswitch Barcodes for Multiplexed Biomarker Profiling. Nano Lett. 2021. https://doi.org/10.1021/acs.nanolett.0c03929.

(21) Chandrasekaran, A. R.; Trivedi, R.; Halvorsen, K. Ribonuclease-Responsive DNA Nanoswitches. Cell Rep. Phys. Sci. 2020, 1 (7), 100117. https://doi.org/10.1016/j.xcrp.2020.100117.

(22) Rothemund, P. W. K. Folding DNA to Create Nanoscale Shapes and Patterns. Nature 2006, 440 (7082), 297–302. https://doi.org/10.1038/nature04586.

(23) Wang, W.-X.; Rajeev, B. W.; Stromberg, A. J.; Ren, N.; Tang, G.; Huang, Q.; Rigoutsos, I.; Nelson, P. T. The Expression of MicroRNA MiR-107 Decreases Early in Alzheimer’s Disease and May Accelerate Disease Progression through Regulation of β-Site Amyloid Precursor Protein-Cleaving Enzyme 1. J. Neurosci. 2008, 28 (5), 1213–1223. https://doi.org/10.1523/JNEUROSCI.5065-07.2008.

(24) Chopra, N.; Wang, R.; Maloney, B.; Nho, K.; Beck, J. S.; Pourshafie, N.; Niculescu, A.; Saykin, A. J.; Rinaldi, C.; Counts, S. E.; Lahiri, D. K. MicroRNA-298 Reduces Levels of Human Amyloid-β Precursor Protein (APP), β-Site APP-Converting Enzyme 1 (BACE1) and Specific Tau Protein Moieties. Mol. Psychiatry 2020, 1–22. https://doi.org/10.1038/s41380-019-0610-2.

(25) Parsi, S.; Smith, P. Y.; Goupil, C.; Dorval, V.; Hébert, S. S. Preclinical Evaluation of MiR-15/107 Family Members as Multifactorial Drug Targets for Alzheimer’s Disease. Mol. Ther. - Nucleic Acids 2015, 4, e256. https://doi.org/10.1038/mtna.2015.33.

(26) Liu, W.; Liu, C.; Zhu, J.; Shu, P.; Yin, B.; Gong, Y.; Qiang, B.; Yuan, J.; Peng, X. MicroRNA-16 Targets Amyloid Precursor Protein to Potentially Modulate Alzheimer’s-Associated Pathogenesis in SAMP8 Mice. Neurobiol. Aging 2012, 33 (3), 522–534. https://doi.org/10.1016/j.neurobiolaging.2010.04.034.

(27) Baker, M. MicroRNA Profiling: Separating Signal from Noise. Nat. Methods 2010, 7 (9), 687–692. https://doi.org/10.1038/nmeth0910-687.

(28) Hao, Y.; Zhao, Y.; Zhao, X.; He, C.; Pang, X.; Wu, T.-C.; Califano, J. A.; Gu, X. Improvement of Prostate Cancer Detection by Integrating the PSA Test With MiRNA Expression Profiling. Cancer Invest. 2011, 29 (4), 318–324. https://doi.org/10.3109/07357907.2011.554477.

(29) Zhao, Y.; Bhattacharjee, S.; Jones, B. M.; Dua, P.; Alexandrov, P. N.; Hill, J. M.; Lukiw, W. J. Regulation of TREM2 Expression by an NF-КB-Sensitive MiRNA-34a. NeuroReport 2013, 24 (6), 318–323. https://doi.org/10.1097/WNR.0b013e32835fb6b0.

(30) Lukiw, W. J. NF-ΚB-Regulated, Proinflammatory MiRNAs in Alzheimer’s Disease. Alzheimers Res. Ther. 2012, 4 (6), 47. https://doi.org/10.1186/alzrt150.

(31) Satoh, J. MicroRNAs and Their Therapeutic Potential for Human Diseases: <BR>Aberrant MicroRNA Expression in Alzheimer’s Disease Brains. J. Pharmacol. Sci. 2010, advpub, 1010080464–1010080464. https://doi.org/10.1254/jphs.10R11FM.

(32) Nagaraj, S.; Zoltowska, K. M.; Laskowska-Kaszub, K.; Wojda, U. MicroRNA Diagnostic Panel for Alzheimer’s Disease and Epigenetic Trade-off between Neurodegeneration and Cancer. Ageing Res. Rev. 2019, 49, 125–143. https://doi.org/10.1016/j.arr.2018.10.008.

(33) Kumar, S.; Reddy, P. H. A New Discovery of MicroRNA-455-3p in Alzheimer’s Disease. J. Alzheimers Dis. 2019, 72 (1), S117–S130. https://doi.org/10.3233/JAD-190583.

(34) Lee, Y. S.; Kim, H. K.; Chung, S.; Kim, K.-S.; Dutta, A. Depletion of Human Micro-RNA MiR-125b Reveals That It Is Critical for the Proliferation of Differentiated Cells but Not for the Down-Regulation of Putative Targets during Differentiation. J. Biol. Chem. 2005, 280 (17), 16635– 16641. https://doi.org/10.1074/jbc.M412247200.

(35) Hébert, S. S.; Horré, K.; Nicolaï, L.; Papadopoulou, A. S.; Mandemakers, W.; Silahtaroglu, A. N.; Kauppinen, S.; Delacourte, A.; Strooper, B. D. Loss of MicroRNA Cluster MiR-29a/b-1 in Sporadic Alzheimer’s Disease Correlates with Increased BACE1/β-Secretase Expression. Proc. Natl. Acad. Sci. 2008, 105 (17), 6415–6420. https://doi.org/10.1073/pnas.0710263105.

(36) Geekiyanage, H.; Chan, C. MicroRNA-137/181c Regulates Serine Palmitoyltransferase and In Turn Amyloid β, Novel Targets in Sporadic Alzheimer’s Disease. J. Neurosci. 2011, 31 (41), 14820–14830. https://doi.org/10.1523/JNEUROSCI.3883-11.2011.

(37) Chandrasekaran, A. R.; Levchenko, O.; Patel, D. S.; MacIsaac, M.; Halvorsen, K. Addressable Configurations of DNA Nanostructures for Rewritable Memory. Nucleic Acids Res. 2017, 45 (19), 11459–11465. https://doi.org/10.1093/nar/gkx777.

(38) Schwarzenbach, H.; da Silva, A. M.; Calin, G.; Pantel, K. Data Normalization Strategies for MicroRNA Quantification. Clin. Chem. 2015, 61 (11), 1333–1342. https://doi.org/10.1373/clinchem.2015.239459.

(39) Noelker, C.; Hampel, H.; Dodel, R. Blood-Based Protein Biomarkers for Diagnosis and Classification of Neurodegenerative Diseases. Mol. Diagn. Ther. 2011, 15 (2), 83–102. https://doi.org/10.1007/BF03256398.

(40) Kawikova, I.; Askenase, P. W. Diagnostic and Therapeutic Potentials of Exosomes in CNS Diseases. Brain Res. 2015, 1617, 63–71. https://doi.org/10.1016/j.brainres.2014.09.070.

