## Supporting Information for "Detection of Alzheimer’s associated microRNAs using a DNA-based smart reagent"

### MATERIALS AND METHODS

#### Materials

Oligonucleotides were purchased from Integrated DNA Technologies (IDT) with standard desalting. M13 circular DNA and BtsCI enzymes were purchased from New England Biolabs (NEB). GelRed nucleic acid stain was purchased from Biotium, Fremont, CA, USA. Molecular biology grade agarose was purchased from Fisher BioReagents. Healthy and Alzheimer's disease brain total RNA was purchased from BioChain Institute, Inc.

#### Linearization of M13 DNA

5 µl of 100 nM circular single-stranded M13 DNA, 2.5 µl of 10× Cut Smart buffer, 0.5 µl of 100 µM BtsCI restriction-site complementary-oligonucleotide and 16 µl of deionized water were mixed and annealed from 95 °C to 50 °C in a T100™ Thermal Cycler (Bio-Rad, Hercules, CA, USA). 1 µl of the BtsCI enzyme (20,000 units/ml) was added to the mixture and incubated at 50 °C for 15 min. The mixture was brought up to 95 °C for 1 min to heat deactivate the enzyme followed by cooling down to 4 °C.

#### Construction of nanoswitches

Linearized single-stranded M13 DNA (20 nM) was mixed with 10-fold excess of the backbone oligonucleotides and detector strands and annealed from 90 °C to 20 °C at 1 °C $min^{-1}$ in a thermal cycler.[1] Following construction, the nanoswitches were purified using UHPLC[2] to remove excess oligonucleotides or used unpurified after dilution in 1× PBS to a concentration of 400 pM.

#### MicroRNA detection (synthetic)

In general, assays were performed by incubating the purified nanoswitches (~100 pM) with the target solution containing microRNA. Concentration measurements were determined by measuring $A_{260}$ absorbance with a Nanodrop spectrophotometer (Thermo Scientific) and applying either molar extinction coefficients (for oligos) or using the standard ssRNA setting (for RNA extracts). Concentrations of final solutions were determined by the volume ratios during dilutions. Typical reactions were carried out in PCR tubes with 10 µl final volumes. Specific protocol variations for each experiment are detailed as follows.

For sensitivity experiment, the target microRNAs were spiked into a 500 nM solution of off-target "blocking" oligos to minimize loss to the tubes. Microcentrifuge tubes and pipette tips were additionally pre-incubated in blocking oligo solution to minimize loss of the target microRNA. The nanoswitch and microRNA solution were incubated overnight at room temperature in a solution containing nominally 1x PBS and 10 mM $MgCl_2$. GelRed and a Ficoll based loading dye (15% Ficoll, 0.1% bromophenol blue) were added to each sample to get a 1x concentration of each. 10 µl of the samples were loaded and run on an unstained agarose gel. For specificity experiments, the final concentration of target strands were 2.5 nM and incubation time was 1 h at room temperature in a solution containing nominally 1x PBS and 10 mM $MgCl_2$.

For multiplexed detection, the 4 individual nanoswitches were concentration normalized and mixed in equal volumes. Nanoswitches at ~100 pM each were mixed with specific combinations of microRNAs (at 2.5 nM final concentration) in a solution containing 1x PBS and 10 mM $MgCl_2$ and incubated at room temperature for 4 hours. GelRed and loading dye were added after incubation and samples were run in an unstained gel. Gels were run at 4 °C for 3 hours at 55 V, with the longer time facilitating easier recognition of the 4 individual looped bands.

**MicroRNA detection from brain total RNA**

For detection of microRNA from brain total RNA samples, 500 ng of total RNA was mixed with the nanoswitches and incubated in a decreasing temperature ramp from 40 °C to 25 °C with 5 minutes for each 0.1 °C (~12.5 hours total). The solution also contained 1x PBS and 1x GelRed during incubation. Ficoll based loading dye was added to the samples before loading and running in an unstained gel. Gels were run at room temperature for 45 minutes at 75 V. Similar protocol was use for multiplexed detection of miR-29a-3p and miR-29b-3p using a nanoswitch mix containing the two nanoswitches corresponding to miR-29a-3p and miR-29b-3p.

**Gel electrophoresis**

Nanoswitches were run in 0.8% agarose gels, cast from molecular biology grade agarose (Fisher BioReagents) dissolved in 0.5× Tris-borate EDTA (TBE) (Ultra-pure grade, Amresco, Solon, OH, USA). Gels were typically run at 75 V (constant voltage) at room temperature. Samples were pre-stained by mixing 1× GelRed stain (Biotium, Fremont, CA, USA) with the samples before loading (variation for total RNA experiment noted above).

**Gel imaging and analysis**

Imaging was done on a Bio-Rad Gel Doc XR+ imager using the default settings for GelRed with UV illumination. Images were typically taken at multiple exposures ranging from 5 seconds to 45 seconds to facilitate accurate quantification. For each gel band, quantification was done using the highest exposure image that did not contain saturated pixels in the band of interest.[3] To quantify each gel band, 12 bit images were imported into ImageJ and processed with a 2 pixel median filter to help reduce noise and small speckles and eliminate hot pixels. Gel images were then rotated 90 degrees and a rectangular selection (of common size) was applied to each gel lane. The plot profile command was used to get a mean intensity profile along the band width, and the area under the curve was quantified as a measure of overall signal. While this analysis can also be done in ImageJ, we preferred to export the profiles to Origin 2017 (OriginLab) for more precise quantification. In Origin, we used the peak analysis tool to draw a spline interpolated baseline and measure the area between the profile and baseline. For the more complex multiplexing experiment we instead explicitly subtracted the baseline using the same method and fit the subsequent profile with 4 gaussian curves.

**Data analysis**

Fold changes were calculated for microRNAs using mean detection signal values obtained from healthy and AD total RNA samples. Error values were determined as the standard deviation of triplicate measurements, and error values were propagated to calculate errors in fold changes.

**References**

1. Chandrasekaran, A. R.; Dey, B. K.; Halvorsen, K. How to perform miRacles: A step-by-step microRNA detection protocol using DNA nanoswitches. *Curr. Protoc. Mol. Biol.* **2020**, *130* (1), e114.

2. Halvorsen, K.; Kizer, M. E.; Wang, X.; Chandrasekaran, A. R.; Basanta-Sanchez, M. Shear dependent LC purification of an engineered DNA nanoswitch and implications for DNA origami. *Anal. Chem.* **2017**, 89, 5673-5677.

3. Chandrasekaran, A. R.; MacIsaac, M.; Dey, P.; Levchenko, O.; Zhou, L.; Andres, M.; Dey, B. K.; Halvorsen, K. Cellular microRNA detection with miRacles: MicroRNA-activated conditional looping of engineered switches. *Sci. Adv.* **2019**, 5, eaau9443.

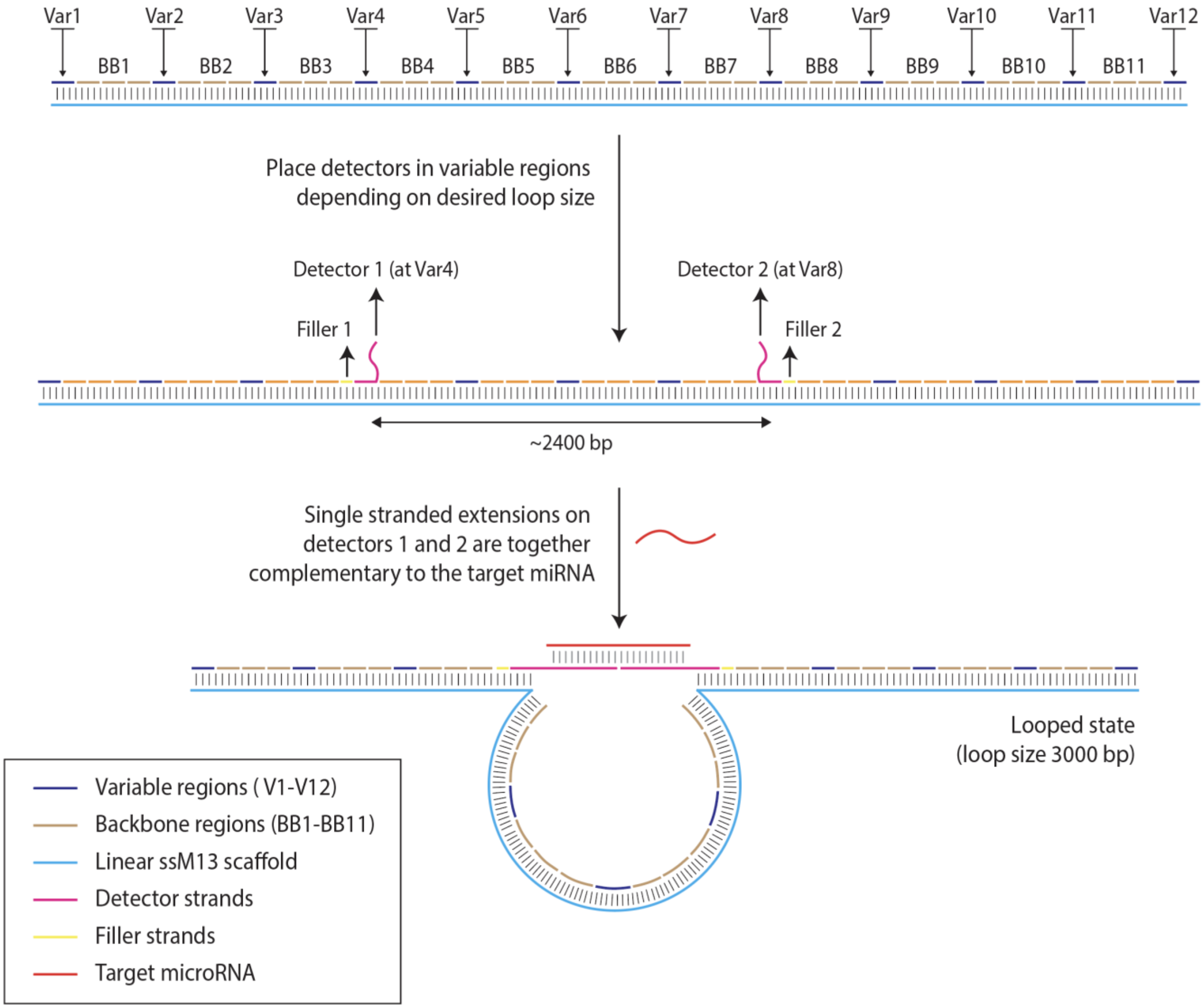

**Figure S1.** *Programmable design of the DNA nanoswitch*. The microRNA detecting nanoswitches are fully double stranded with the exception of the two single-stranded overhangs that complement the microRNA of interest. To accomplish this, the linear M13 (light blue) is hybridized with over 120 oligonucleotides. To facilitate mixing, we have designated 12 regions (60 nt each) as "variable" regions where we can either use fully complementary strands or swap in different detector strands with overhangs. Due to length constraints on oligo synthesis, variable regions that accommodate detector strands often require an additional "filler" strand. In a typical situation, the detector strand may bind 40 of the 60 nt in the variable region, and a filler strand would bind the other 20 nt. Aside from the variable regions, the remainder of the M13 is covered by 109 "backbone" oligos (60 nt each except one 49 nt region). The distance between the two detectors dictates the loop size and migration of the looped state on a gel.

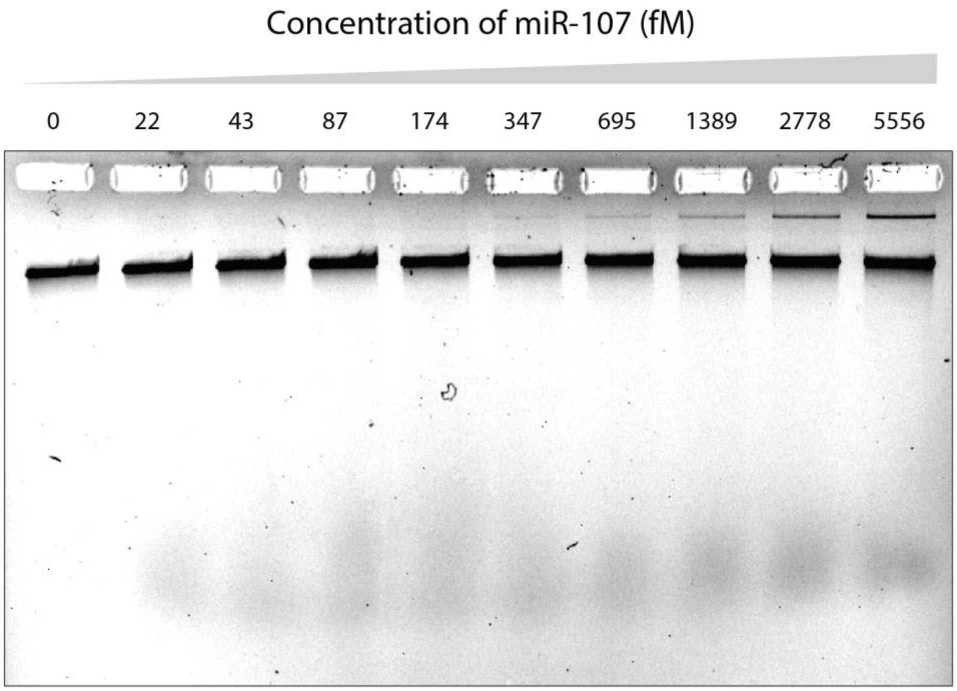

**Figure S2**. Sensitivity of miR-107 (full gel of result shown in Figure 1c).

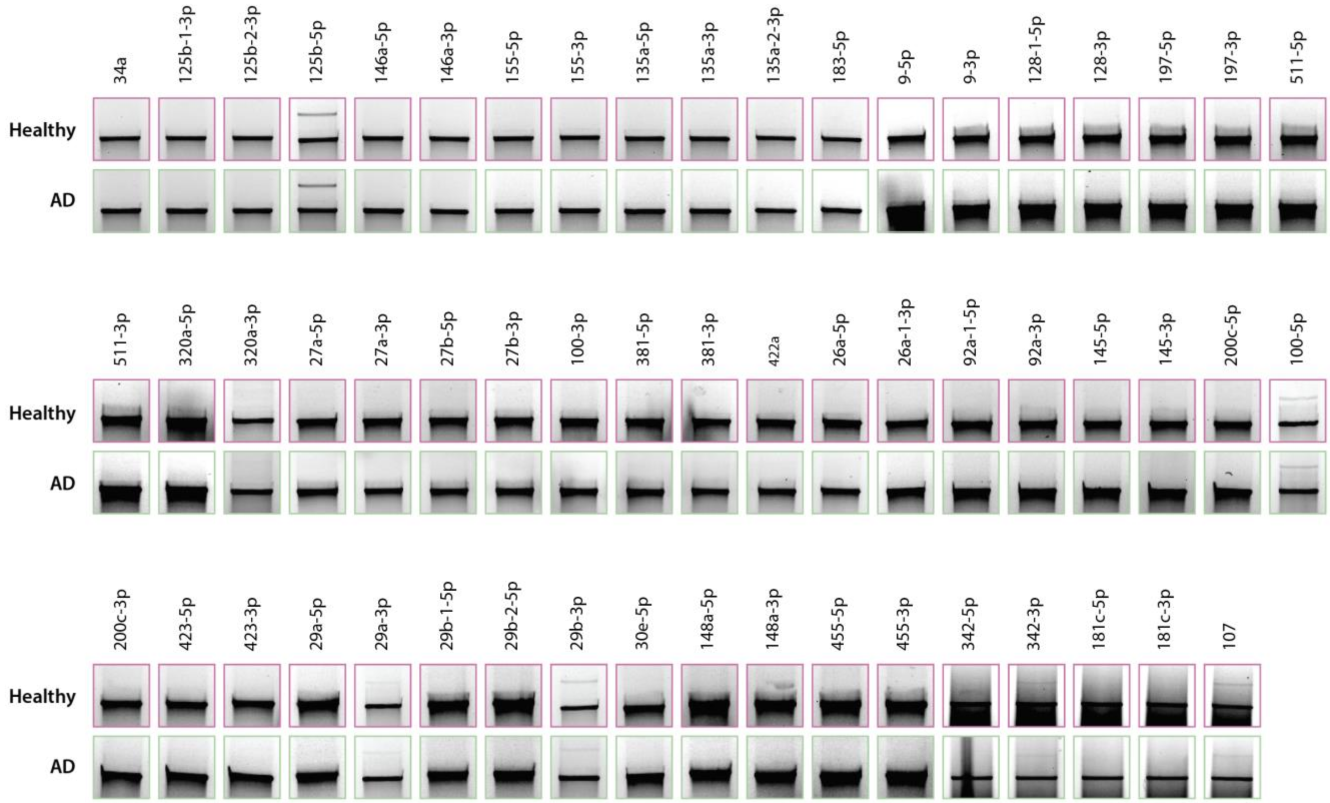

**Figure S3**. Screening of AD-related microRNAs in total RNA extracts from healthy and AD brain total RNA.

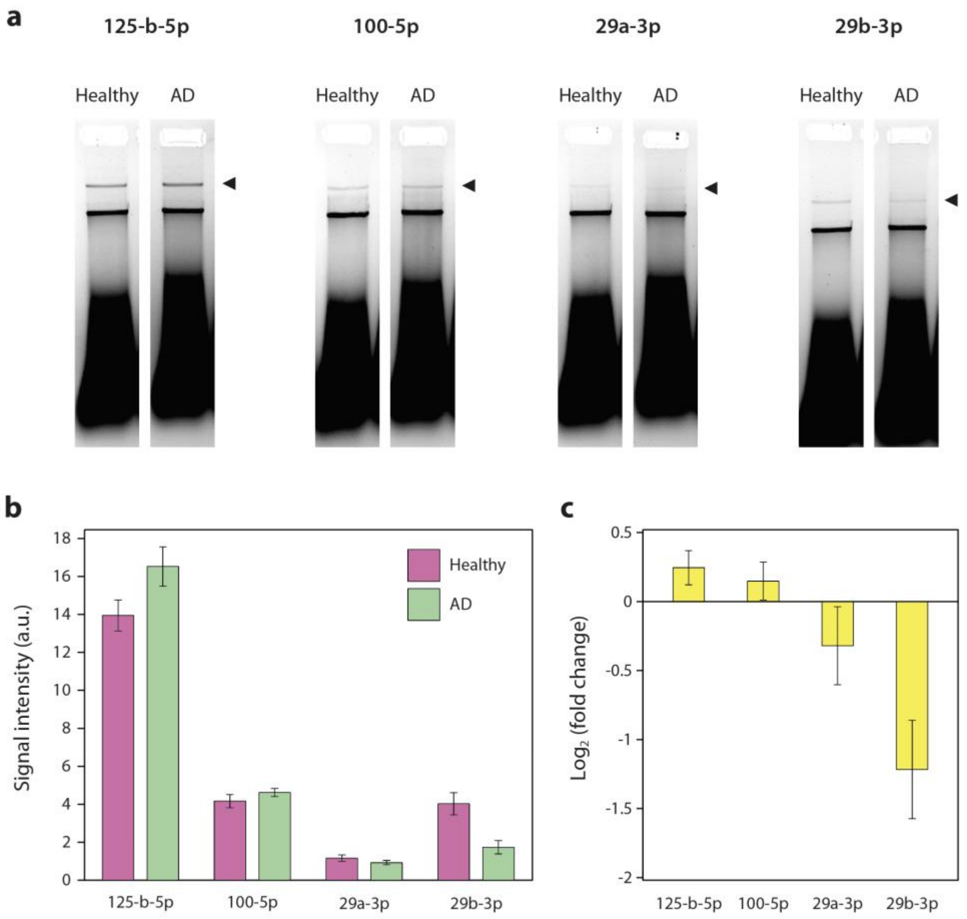

**Figure S4**. (a) Gel images showing detection of selected microRNAs from brain total RNA. (b) Quantified results for microRNA 125-b-5p, 100-5p, 29a-3p and 29b-3p and (c) fold change in expression compared to healthy samples, showing upregulation of miR-125-b-5p and 100-5p, and downregulation of miR-29a-3p and 29-b-3p. Healthy and AD brain total RNA samples with nanoswitches were run in the same gel with replicates and analyzed. Gel lanes are sliced and placed next to each other for visual clarity.

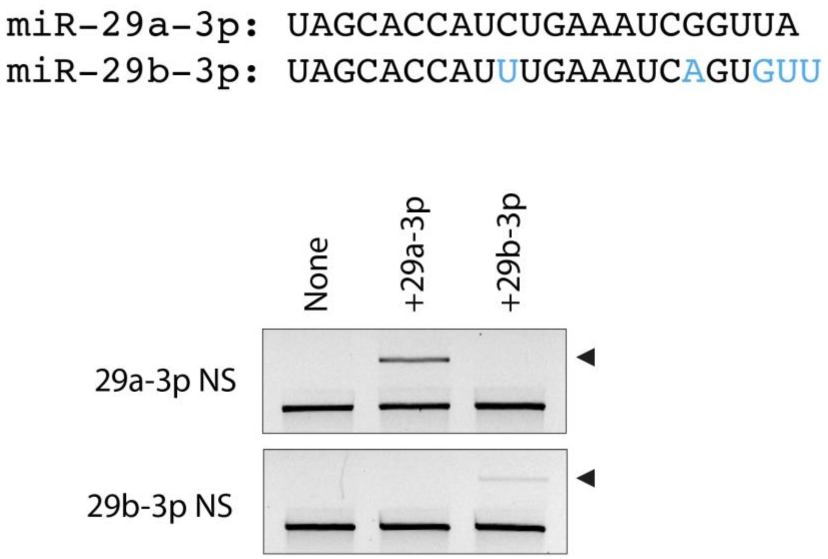

**Figure S5**. Gels showing specific detection of miR-29a-3p and miR-29b-3p.

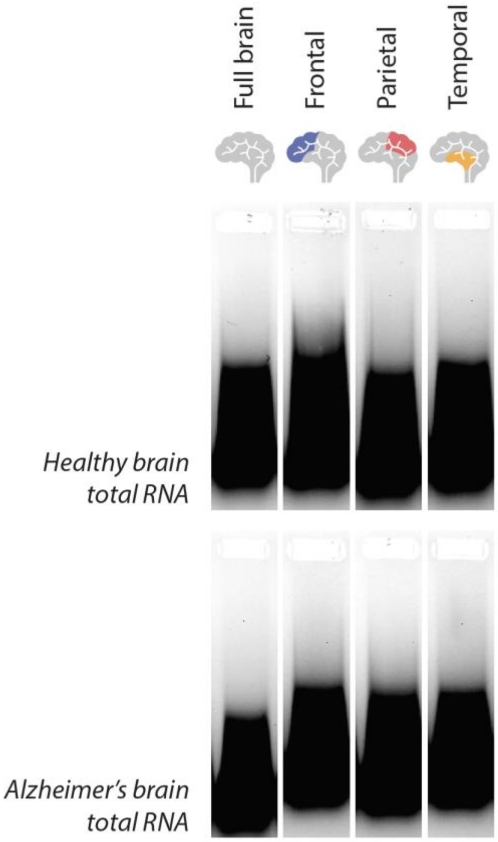

**Figure S6**. Control gel showing total RNA from full brain, frontal, parietal, and temporal regions of both healthy and AD brain samples. There are no bands in the total RNA that migrate similar to linear or looped DNA nanoswitches.

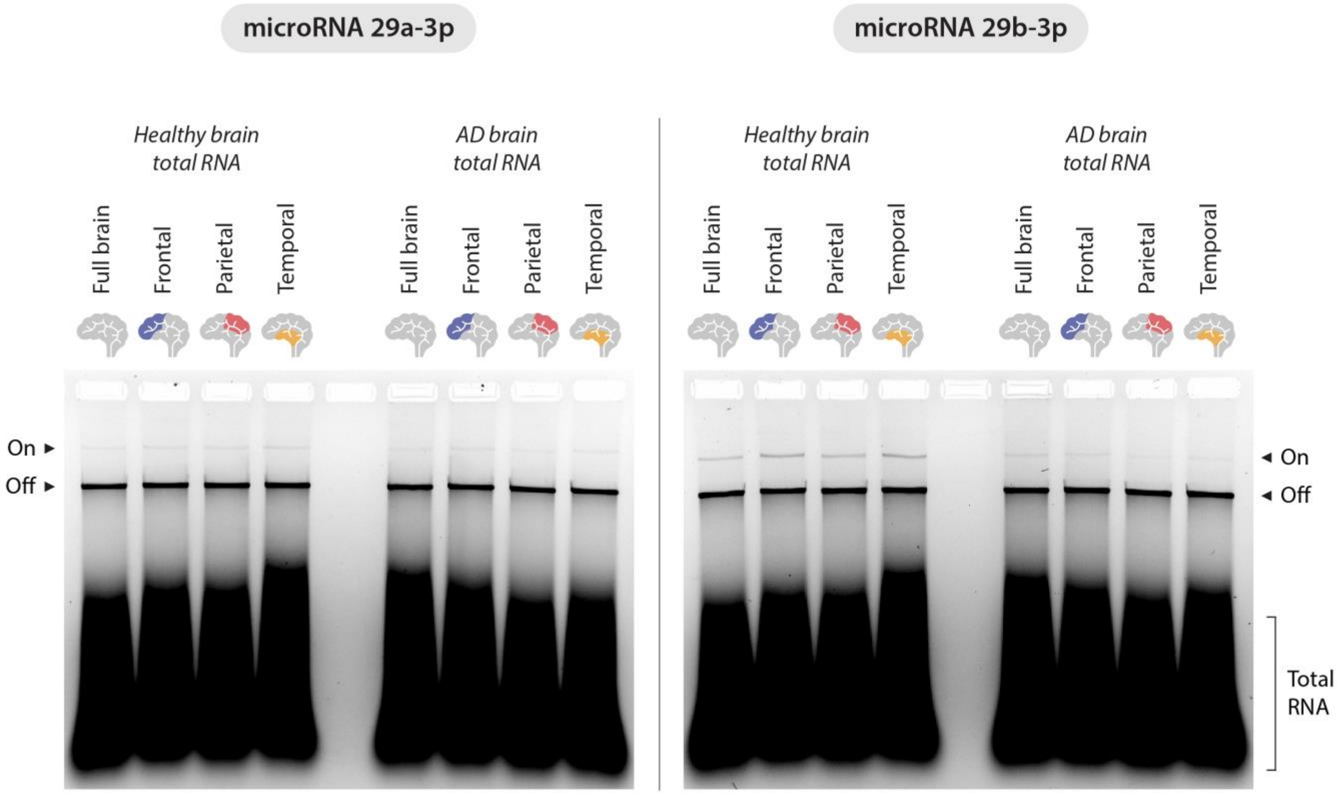

**Figure S7**. Gels showing detection of miR-29a-3p and miR-29b-3p from healthy and AD total RNA extracts (full images of gels shown in figure 2c, 2f).

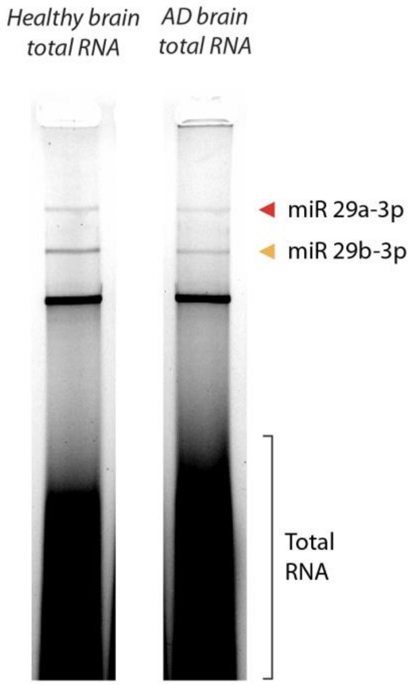

**Figure S8**. Multiplexed detection of miR-29a-3p and miR-29b-3p in a single assay (full gels of images shown in figure 3a).

| Material | Unit cost | Amount needed/lane | Cost per reaction | Vendor |
|---|---|---|---|---|
| ***DNA and enzymes*** | | | | |
| Scaffold DNA | $ 3.40 /µg | 0.0024 µg | $ 0.008 | NEB |
| Oligonucleotides | $ 0.37 /µg | 0.0024 µg | $ 0.001 | IDT |
| Linearization enzyme | $ 0.68 /µL | 0.0032 µL | $ 0.002 | NEB |
| ***Gel reagents*** | | | | |
| Agarose | $ 0.84 /g | 0.02 g | $ 0.01680 | Sigma |
| GelRed | $ 0.23 /µL | 0.0001 µL | $ 0.00002 | Biotium |
| Ficoll (loading dye) | $ 0.21 /g | 0.0003 g | $ 0.00006 | VWR |
| ***Total*** | | | $ 0.02788 | |

**Table S1.** Cost per reaction based on 1000 amol of nanoswitch per reaction.

Complete list of all sequences used. All sequences are written from 5' to 3'.

| Backbone oligonucleotides | | |
|---|---|---|
| **BB#** | **Sequence** | **Length** |
| 1 | AGAGCATAAAGCTAAATCGGTTGTACCAAAAACATTATGACCCTGTAATACTTTTGCGGG | 60 |
| 2 | AGAAGCCTTTATTTCAACGCAAGGATAAAAATTTTTAGAACCCTCATATATTTTAAATGC | 60 |
| 3 | AATGCCTGAGTAATGTGTAGGTAAAGATTCAAAAGGGTGAGAAAGGCCGGAGACAGTCAA | 60 |
| 4 | ATCACCATCAATATGATATTCAACCGTTCTAGCTGATAAATTAATGCCGGAGAGGGTAGC | 60 |
| 5 | TATTTTTGAGAGATCTACAAAGGCTATCAGGTCATTGCCTGAGAGTCTGGAGCAAACAAG | 60 |
| 6 | AGAATCGATGAACGGTAATCGTAAAACTAGCATGTCAATCATATGTACCCCGGTTGATAA | 60 |
| 7 | TCAGAAAAGCCCCAAAAACAGGAAGATTGTATAAGCAAATATTTAAATTGTAAACGTTAA | 60 |
| 8 | TATTTTGTTAAAATTCGCATTAAATTTTTGTTAAATCAGCTCATTTTTTAACCAATAGGA | 60 |
| 9 | ACGCCATCAAAAATAATTCGCGTCTGGCCTTCCTGTAGCCAGCTTTCATCAACATTAAAT | 60 |
| 10 | GGATAGGTCACGTTGGTGTAGATGGGCGCATCGTAACCGTGCATCTGCCAGTTTGAGGGG | 60 |
| 11 | ACGACGACAGTATCGGCCTCAGGAAGATCGCACTCCAGCCAGCTTTCCGGCACCGCTTCT | 60 |
| 12 | GGTGCCGGAAACCAGGCAAAGCGCCATTCGCCATTCAGGCTGCGCAACTGTTGGGAAGGG | 60 |
| 13 | CGATCGGTGCGGGCCTCTTCGCTATTACGCCAGCTGGCGAAAGGGGGATGTGCTGCAAGG | 60 |
| 14 | CGATTAAGTTGGGTAACGCCAGGGTTTTCCCAGTCACGACGTTGTAAAACGACGGCCAGT | 60 |
| 15 | GCCAAGCTTGCATGCCTGCAGGTCGACTCTAGAGGATCCCCGGGTACCGAGCTCGAATTC | 60 |
| 16 | GTAATCATGGTCATAGCTGTTTCCTGTGTGAAATTGTTATCCGCTCACAATTCCACACAA | 60 |
| 17 | CATACGAGCCGGAAGCATAAAGTGTAAAGCCTGGGGTGCCTAATGAGTGAGCTAACTCAC | 60 |
| 18 | ATTAATTGCGTTGCGCTCACTGCCCGCTTTCCAGTCGGGAAACCTGTCGTGCCAGCTGCA | 60 |
| 19 | TTAATGAATCGGCCAACGCGCGGGGAGAGGCGGTTTGCGTATTGGGCGCCAGGGTGGTTT | 60 |
| 20 | GTTGCAGCAAGCGGTCCACGCTGGTTTGCCCCAGCAGGCGAAAATCCTGTTTGATGGTGG | 60 |
| 21 | TTCCGAAATCGGCAAAATCCCTTATAAATCAAAAGAATAGCCCGAGATAGGGTTGAGTGT | 60 |
| 22 | TGTTCCAGTTTGGAACAAGAGTCCACTATTAAAGAACGTGGACTCCAACGTCAAAGGGCG | 60 |
| 23 | AAAAACCGTCTATCAGGGCGATGGCCCACTACGTGAACCATCACCCAAATCAAGTTTTTT | 60 |
| 24 | GGGGTCGAGGTGCCGTAAAGCACTAAATCGGAACCCTAAAGGGAGCCCCCGATTTAGAGC | 60 |
| 25 | TTGACGGGGAAAGCCGGCGAACGTGGCGAGAAAGGAAGGGAAGAAAGCGAAAGGAGCGGG | 60 |
| 26 | CGCTAGGGCGCTGGCAAGTGTAGCGGTCACGCTGCGCGTAACCACCACACCCGCCGCGCT | 60 |
| 27 | TAATGCGCCGCTACAGGGCGCGTACTATGGTTGCTTTGACGAGCACGTATAACGTGCTTT | 60 |
| 28 | CCTCGTTAGAATCAGAGCGGGAGCTAAACAGGAGGCCGATTAAAGGGATTTTAGACAGGA | 60 |
| 29 | ACGGTACGCCAGAATCCTGAGAAGTGTTTTTATAATCAGTGAGGCCACCGAGTAAAAGAG | 60 |
| 30 | TTGCCTGAGTAGAAGAACTCAAACTATCGGCCTTGCTGGTAATATCCAGAACAATATTAC | 60 |
| 31 | CGCCAGCCATTGCAACAGGAAAAACGCTCATGGAAATACCTACATTTTGACGCTCAATCG | 60 |
| 32 | TCTGAAATGGATTATTTACATTGGCAGATTCACCAGTCACACGACCAGTAATAAAAGGGA | 60 |
| 33 | CATTCTGGCCAACAGAGATAGAACCCTTCTGACCTGAAAGCGTAAGAATACGTGGCACAG | 60 |
| 34 | ACAATATTTTTGAATGGCTATTAGTCTTTAATGCGCGAACTGATAGCCCTAAAACATCGC | 60 |
| 35 | CATTAAAAATACCGAACGAACCACCAGCAGAAGATAAAACAGAGGTGAGGCGGTCAGTAT | 60 |
| 36 | TAACACCGCCTGCAACAGTGCCACGCTGAGAGCCAGCAGCAAATGAAAAATCTAAAGCAT | 60 |
| 37 | CACCTTGCTGAACCTCAAATATCAAACCCTCAATCAATATCTGGTCAGTTGGCAAATCAA | 60 |
| 38 | CAGTTGAAAGGAATTGAGGAAGGTTATCTAAAATATCTTTAGGAGCACTAACAACTAATA | 60 |

| | | |
|---|---|---|
| 39 | GATTAGAGCCGTCAATAGATAATACATTTGAGGATTTAGAAGTATTAGACTTTACAAACA | 60 |
| 40 | CATTATCATTTTGCGGAACAAAGAAACCACCAGAAGGAGCGGAATTATCATCATATTCCT | 60 |
| 41 | GATTATCAGATGATGGCAATTCATCAATATAATCCTGATTGTTTGGATTATACTTCTGAA | 60 |
| 42 | TAATGGAAGGGTTAGAACCTACCATATCAAAATTATTTGCACGTAAAACAGAAATAAAGA | 60 |
| 43 | AATTGCGTAGATTTTCAGGTTTAACGTCAGATGAATATACAGTAACAGTACCTTTTACAT | 60 |
| 44 | CGGGAGAAACAATAACGGATTCGCCTGATTGCTTTGAATACCAAGTTACAAAATCGCGCA | 60 |
| 45 | GAGGCGAATTATTCATTTCAATTACCTGAGCAAAAGAAGATGATGAAACAAACATCAAGA | 60 |
| 46 | AAACAAAATTAATTACATTTAACAATTTCATTTGAATTACCTTTTTTAATGGAAACAGTA | 60 |
| 47 | CATAAATCAATATATGTGAGTGAATAACCTTGCTTCTGTAAATCGTCGCTATTAATTAAT | 60 |
| 48 | TTTCCCTTAGAATCCTTGAAAACATAGCGATAGCTTAGATTAAGACGCTGAGAAGAGTCA | 60 |
| 49 | ATAGTGAATTTATCAAAATCATAGGTCTGAGAGACTACCTTTTTAACCTCCGGCTTAGGT | 60 |
| 50 | GAAAACTTTTTCAAATATATTTTAGTTAATTTCATCTTCTGACCTAAATTTAATGGTTTG | 60 |
| 51 | AAATACCGACCGTGTGATAAATAAGGCGTTAAATAAGAATAAACACCGGAATCATAATTA | 60 |
| 52 | CTAGAAAAAGCCTGTTTAGTATCATATGCGTTATACAAATTCTTACCAGTATAAAGCCAA | 60 |
| 53 | CGCTCAACAGTAGGGCTTAATTGAGAATCGCCATATTTAACAACGCCAACATGTAATTTA | 60 |
| 54 | GGCAGAGGCATTTTCGAGCCAGTAATAAGAGAATATAAAGTACCGACAAAAGGTAAAGTA | 60 |
| 55 | ATTCTGTCCAGACGACGACAATAAACAACATGTTCAGCTAATGCAGAACGCGCCTGTTTA | 60 |
| 56 | TCAACAATAGATAAGTCCTGAACAAGAAAAATAATATCCCATCCTAATTTACGAGCATGT | 60 |
| 57 | AGAAACCAATCAATAATCGGCTGTCTTTCCTTATCATTCCAAGAACGGGTATTAAACCAA | 60 |
| 58 | GTACCGCACTCATCGAGAACAAGCAAGCCGTTTTTATTTTCATCGTAGGAATCATTACCG | 60 |
| 59 | CGCCCAATAGCAAGCAAATCAGATATAGAAGGCTTATCCGGTATTCTAAGAACGCGAGGC | 60 |
| 60 | ATTTTGCACCCAGCTACAATTTTATCCTGAATCTTACCAACGCTAACGAGCGTCTTTCCA | 60 |
| 61 | GAGCCTAATTTGCCAGTTACAAAATAAACAGCCATATTATTTATCCCAATCCAAATAAGA | 60 |
| 62 | AACGATTTTTTGTTTAACGTCAAAAATGAAAATAGCAGCCTTTACAGAGAGAATAACATA | 60 |
| 63 | AAAACAGGGAAGCGCATTAGACGGGAGAATTAACTGAACACCCTGAACAAAGTCAGAGGG | 60 |
| 64 | TAATTGAGCGCTAATATCAGAGAGATAACCCACAAGAATTGAGTTAAGCCCAATAATAAG | 60 |
| 65 | AGCAAGAAACAATGAAATAGCAATAGCTATCTTACCGAAGCCCTTTTTAAGAAAAGTAAG | 60 |
| 66 | CAGATAGCCGAACAAAGTTACCAGAAGGAAACCGAGGAAACGCAATAATAACGGAATACC | 60 |
| 67 | CAAAAGAACTGGCATGATTAAGACTCCTTATTACGCAGTATGTTAGCAAACGTAGAAAAT | 60 |
| 68 | ACATACATAAAGGTGGCAACATATAAAAGAAACGCAAAGACACCACGGAATAAGTTTATT | 60 |
| 69 | TTGTCACAATCAATAGAAAATTCATATGGTTTACCAGCGCCAAAGACAAAAGGGCGACAT | 60 |
| 70 | TCACCGTCACCGACTTGAGCCATTTGGGAATTAGAGCCAGCAAAATCACCAGTAGCACCA | 60 |
| 71 | TTACCATTAGCAAGGCCGGAAACGTCACCAATGAAACCATCGATAGCAGCACCGTAATCA | 60 |
| 72 | GTAGCGACAGAATCAAGTTTGCCTTTAGCGTCAGACTGTAGCGCGTTTTCATCGGCATTT | 60 |
| 73 | TCGGTCATAGCCCCCTTATTAGCGTTTGCCATCTTTTCATAATCAAAATCACCGGAACCA | 60 |
| 74 | GAGCCACCACCGGAACCGCCTCCCTCAGAGCCGCCACCCTCAGAACCGCCACCCTCAGAG | 60 |
| 75 | CCACCACCCTCAGAGCCGCCACCAGAACCACCACCAGAGCCGCCGCCAGCATTGACAGGA | 60 |
| 76 | GGTTGAGGCAGGTCAGACGATTGGCCTTGATATTCACAAACAAATAAATCCTCATTAAAG | 60 |
| 77 | CCAGAATGGAAAGCGCAGTCTCTGAATTTACCGTTCCAGTAAGCGTCATACATGGCTTTT | 60 |
| 78 | GATGATACAGGAGTGTACTGGTAATAAGTTTTAACGGGGTCAGTGCCTTGAGTAACAGTG | 60 |
| 79 | CCCGTATAAACAGTTAATGCCCCCTGCCTATTTCGGAACCTATTATTCTGAAACATGAAA | 60 |
| 80 | CCAGGCGGATAAGTGCCGTCGAGAGGGTTGATATAAGTATAGCCCGGAATAGGTGTATCA | 60 |
| 81 | CCGTACTCAGGAGGTTTAGTACCGCCACCCTCAGAACCGCCACCCTCAGAACCGCCACCC | 60 |

| | | |
|---|---|---|
| 82 | TCAGAGCCACCACCCTCATTTTCAGGGATAGCAAGCCCAATAGGAACCCATGTACCGTAA | 60 |
| 83 | CACTGAGTTTCGTCACCAGTACAAACTACAACGCCTGTAGCATTCCACAGACAGCCCTCA | 60 |
| 84 | TAGTTAGCGTAACGATCTAAAGTTTTGTCGTCTTTCCAGACGTTAGTAAATGAATTTTCT | 60 |
| 85 | GTATGGGATTTTGCTAAACAACTTTCAACAGTTTCAGCGGAGTGAGAATAGAAAGGAACA | 60 |
| 86 | ACTAAAGGAATTGCGAATAATAATTTTTTCACGTTGAAAATCTCCAAAAAAAAGGCTCCA | 60 |
| 87 | AAAGGAGCCTTTAATTGTATCGGTTTATCAGCTTGCTTTCGAGGTGAATTTCTTAAACAG | 60 |
| 88 | CTTGATACCGATAGTTGCGCCGACAATGACAACAACCATCGCCCACGCATAACCGATATA | 60 |
| 89 | TTCGGTCGCTGAGGCTTGCAGGGAGTTAAAGGCCGCTTTTGCGGGATCGTCACCCTCAGC | 60 |
| 90 | CTTTTTCATGAGGAAGTTTCCATTAAACGGGTAAAATACGTAATGCCACTACGAAGGCAC | 60 |
| 91 | CAACCTAAAACGAAAGAGGCAAAAGAATACACTAAAACACTCATCTTTGACCCCCAGCGA | 60 |
| 92 | TTATACCAAGCGCGAAACAAAGTACAACGGAGATTTGTATCATCGCCTGATAAATTGTGT | 60 |
| 93 | CGAAATCCGCGACCTGCTCCATGTTACTTAGCCGGAACGAGGCGCAGACGGTCAATCATA | 60 |
| 94 | AGGGAACCGAACTGACCAACTTTGAAAGAGGACAGATGAACGGTGTACAGACCAGGCGCA | 60 |
| 95 | TAGGCTGGCTGACCTTCATCAAGAGTAATCTTGACAAGAACCGGATATTCATTACCCAAA | 60 |
| 96 | TCAACGTAACAAAGCTGCTCATTCAGTGAATAAGGCTTGCCCTGACGAGAAACACCAGAA | 60 |
| 97 | CGAGTAGTAAATTGGGCTTGAGATGGTTTAATTTCAACTTTAATCATTGTGAATTACCTT | 60 |
| 98 | ATGCGATTTTAAGAACTGGCTCATTATACCAGTCAGGACGTTGGGAAGAAAAATCTACGT | 60 |
| 99 | TAATAAAACGAACTAACGGAACAACATTATTACAGGTAGAAAGATTCATCAGTTGAGATT | 60 |
| 100 | TAAGAGCAACACTATCATAACCCTCGTTTACCAGACGACGATAAAAACCAAAATAGCGAG | 60 |
| 101 | AGGCTTTTGCAAAAGAAGTTTTGCCAGAGGGGGTAATAGTAAAATGTTTAGACTGGATAG | 60 |
| 102 | CGTCCAATACTGCGGAATCGTCATAAATATTCATTGAATCCCCCTCAAATGCTTTAAACA | 60 |
| 103 | GTTCAGAAAACGAGAATGACCATAAATCAAAAATCAGGTCTTTACCCTGACTATTATAGT | 60 |
| 104 | CAGAAGCAAAGCGGATTGCATCAAAAAGATTAAGAGGAAGCCCGAAAGACTTCAAATATC | 60 |
| 105 | GCGTTTTAATTCGAGCTTCAAAGCGAACCAGACCGGAAGCAAACTCCAACAGGTCAGGAT | 60 |
| 106 | TAGAGAGTACCTTTAATTGCTCCTTTTGATAAGAGGTCATTTTTGCGGATGGCTTAGAGC | 60 |
| 107 | TTAATTGCTGAATATAATGCTGTAGCTCAACATGTTTTAAATATGCAACTAAAGTACGGT | 60 |
| 108 | GTCTGGAAGTTTCATTCCATATAACAGTTGATTCCCAATTCTGCGAACGAGTAGATTTAG | 60 |
| 109 | TTTGACCATTAGATACATTTCGCAAATGGTCAATAACCTGTTTAGCTAT | 49 |

Regions in **green** indicate locations where detector strands are placed. Regions in **blue** are single stranded extensions on detectors that are complementary to two halves of the target microRNA.

| **Variable sequences** | | |
|---|---|---|
| **#** | **Sequence** | **Length** |
| V1 | AACATCCAATAAATCATACAGGCAAGGCAAAGAATTAGCAAAATTAAGCAATAAAGCCTC | 60 |
| V2 | GTGAGCGAGTAACAACCCGTCGGATTCTCCGTGGGAACAAACGGCGGATTGACCGTAATG | 60 |
| V3 | TTCTTTTCACCAGTGAGACGGGCAACAGCTGATTGCCCTTCACCGCCTGGCCCTGAGAGA | 60 |
| V4 | TCTGTCCATCACGCAAATTAACCGTTGTAGCAATACTTCTTTGATTAGTAATAACATCAC | 60 |
| V5 | ATTCGACAACTCGTATTAAATCCTTTGCCCGAACGTTATTAATTTTAAAAGTTTGAGTAA | 60 |
| V6 | TGGGTTATATAACTATATGTAAATGCTGATGCAAATCCAATCGCAAGACAAAGAACGCGA | 60 |
| V7 | GTTTTAGCGAACCTCCCGACTTGCGGGAGGTTTTGAAGCCTTAAATCAAGATTAGTTGCT | 60 |
| V8 | TCAACCGATTGAGGGAGGGAAGGTAAATATTGACGGAAATTATTCATTAAAGGTGAATTA | 60 |
| V9 | GTATTAAGAGGCTGAGACTCCTCAAGAGAAGGATTAGGATTAGCGGGGTTTTGCTCAGTA | 60 |
| V10 | AGCGAAAGACAGCATCGGAACGAGGGTAGCAACGGCTACAGAGGCTTTGAGGACTAAAGA | 60 |
| V11 | TAGGAATACCACATTCAACTAATGCAGATACATAACGCCAAAAGGAATTACGAGGCATAG | 60 |
| V12 | ATTTTCATTTGGGGCGCGAGCTGAAAAGGTGGCATCAATTCTACTAATAGTAGTAGCATT | 60 |

| **MicroRNA detector sequences** | | |
|---|---|---|
| **#** | **Sequence** | **Length** |
| V4 miR-107 | ACCGTTGTAGCAATACTTCTTTGATTAGTAATAACATCACTGATAGCCCTGT | 52 |
| V8 miR-107 | ACAATGCTGCTTCAACCGATTGAGGGAGGGAAGGTAAATATTGACGGAAAT | 51 |
| V4 miR-15a | ACCGTTGTAGCAATACTTCTTTGATTAGTAATAACATCACCACAAACCATT | 51 |
| V8 miR-15a/b | ATGTGCTGCTATCAACCGATTGAGGGAGGGAAGGTAAATATTGACGGAAAT | 51 |
| V4 miR-15b | ACCGTTGTAGCAATACTTCTTTGATTAGTAATAACATCACTGTAAACCATG | 51 |
| V4 miR-16 | ACCGTTGTAGCAATACTTCTTTGATTAGTAATAACATCACCGCCAATATTT | 51 |
| V8 miR-16 | ACGTGCTGCTATCAACCGATTGAGGGAGGGAAGGTAAATATTGACGGAAAT | 51 |
| V4 miR-34a-5p 40-11 | ACCGTTGTAGCAATACTTCTTTGATTAGTAATAACATCACACAACCAGCTA | 51 |
| V8 miR-34a-5p 11-40 | AGACACTGCCATCAACCGATTGAGGGAGGGAAGGTAAATATTGACGGAAAT | 51 |
| V4 miR-125b-1-3p 40-11 | ACCGTTGTAGCAATACTTCTTTGATTAGTAATAACATCACAGCTCCCAAGA | 51 |
| V8 miR-125b-1-3p 11-40 | GCCTAACCCGTTCAACCGATTGAGGGAGGGAAGGTAAATATTGACGGAAAT | 51 |
| V4 miR-125b-2-3p 40-11 | ACCGTTGTAGCAATACTTCTTTGATTAGTAATAACATCACGTCCCAAGAGC | 51 |
| V8 miR-125b-2-3p 11-40 | CTGACTTGTGATCAACCGATTGAGGGAGGGAAGGTAAATATTGACGGAAAT | 51 |
| V4 miR-146a-5p 40-11 | ACCGTTGTAGCAATACTTCTTTGATTAGTAATAACATCACAACCCATGGAA | 51 |
| V8 miR-146a-5p 11-40 | TTCAGTTCTCATCAACCGATTGAGGGAGGGAAGGTAAATATTGACGGAAAT | 51 |
| V4 miR-146a-3p 40-11 | ACCGTTGTAGCAATACTTCTTTGATTAGTAATAACATCACCTGAAGAACTG | 51 |
| V8 miR-146a-3p 10-40 | AATTTCAGAGGTCAACCGATTGAGGGAGGGAAGGTAAATATTGACGGAAAT | 51 |
| V4 miR-155-5p 40-12 | ACCGTTGTAGCAATACTTCTTTGATTAGTAATAACATCACAACCCCTATCAC | 52 |
| V8 miR-155-5p 12-40 | GATTAGCATTAATCAACCGATTGAGGGAGGGAAGGTAAATATTGACGGAAAT | 52 |
| V4 miR-155-3p 40-11 | ACCGTTGTAGCAATACTTCTTTGATTAGTAATAACATCACTGTTAATGCTA | 51 |

| | | |
|---|---|---|
| V8 miR-155-3p 11-40 | ATATGTAGGAGTCAACCGATTGAGGGAGGGAAGGTAAATATTGACGGAAAT | 51 |
| V4 miR-183-5p 40-11 | ACCGTTGTAGCAATACTTCTTTGATTAGTAATAACATCACAGTGAATTCTA | 51 |
| V8 miR-183-5p 11-40 | CCAGTGCCATATCAACCGATTGAGGGAGGGAAGGTAAATATTGACGGAAAT | 51 |
| V4 miR-135a-5p 40-12 | ACCGTTGTAGCAATACTTCTTTGATTAGTAATAACATCACTCACATAGGAAT | 52 |
| V8 miR-135a-5p 11-40 | AAAAAGCCATATCAACCGATTGAGGGAGGGAAGGTAAATATTGACGGAAAT | 51 |
| V4 miR-135a-3p 40-11 | ACCGTTGTAGCAATACTTCTTTGATTAGTAATAACATCACCGCCACGGCTC | 51 |
| V8 miR-135a-3p 11-40 | CAATCCCTATATCAACCGATTGAGGGAGGGAAGGTAAATATTGACGGAAAT | 51 |
| V4 miR-135a-2-3p 40-11 | ACCGTTGTAGCAATACTTCTTTGATTAGTAATAACATCACTTCATGGCTTC | 51 |
| V8 miR-135a-2-3p 11-40 | CATCCCTACATTCAACCGATTGAGGGAGGGAAGGTAAATATTGACGGAAAT | 51 |
| V4 miR-9-5p 40-12 | ACCGTTGTAGCAATACTTCTTTGATTAGTAATAACATCACACAACCAGCTA | 51 |
| V8 miR-9-5p 11-40 | AGACACTGCCATCAACCGATTGAGGGAGGGAAGGTAAATATTGACGGAAAT | 51 |
| V4 miR-9-3p 40-11 | ACCGTTGTAGCAATACTTCTTTGATTAGTAATAACATCACTCACAAGTTAG | 51 |
| V8 miR-9-3p 11-40 | GGTCTCAGGGATCAACCGATTGAGGGAGGGAAGGTAAATATTGACGGAAAT | 51 |
| V4 miR-128-1-5p 40-12 | ACCGTTGTAGCAATACTTCTTTGATTAGTAATAACATCACAGCTCCCAAGA | 51 |
| V8 miR-128-1-5p 11-40 | GCCTAACCCGTTCAACCGATTGAGGGAGGGAAGGTAAATATTGACGGAAAT | 51 |
| V4 miR-128-3p 40-11 | ACCGTTGTAGCAATACTTCTTTGATTAGTAATAACATCACGTCCCAAGAGC | 51 |
| V8 miR-128-3p 10-40 | CTGACTTGTGATCAACCGATTGAGGGAGGGAAGGTAAATATTGACGGAAAT | 51 |
| V4 miR-197-5p 40-12 | ACCGTTGTAGCAATACTTCTTTGATTAGTAATAACATCACAACCCATGGAA | 51 |
| V8 miR-197-5p 11-40 | TTCAGTTCTCATCAACCGATTGAGGGAGGGAAGGTAAATATTGACGGAAAT | 51 |
| V4 miR-197-3p 40-11 | ACCGTTGTAGCAATACTTCTTTGATTAGTAATAACATCACACAACCAGCTA | 51 |
| V8 miR-197-3p 11-40 | AGACACTGCCATCAACCGATTGAGGGAGGGAAGGTAAATATTGACGGAAAT | 51 |
| V4 miR-511-5p 40-11 | ACCGTTGTAGCAATACTTCTTTGATTAGTAATAACATCACTCACAAGTTAG | 51 |
| V8 miR-511-5p 10-40 | GGTCTCAGGGATCAACCGATTGAGGGAGGGAAGGTAAATATTGACGGAAAT | 51 |
| V4 miR-511-3p 40-10 | ACCGTTGTAGCAATACTTCTTTGATTAGTAATAACATCACACAACCAGCTA | 51 |
| V8 miR-511-3p 10-40 | AGACACTGCCATCAACCGATTGAGGGAGGGAAGGTAAATATTGACGGAAAT | 51 |
| V4 miR-320a-5p 40-11 | ACCGTTGTAGCAATACTTCTTTGATTAGTAATAACATCACTCACAAGTTAG | 51 |
| V8 miR-320a-5p 11-40 | GGTCTCAGGGATCAACCGATTGAGGGAGGGAAGGTAAATATTGACGGAAAT | 51 |
| V4 miR-320a-3p 40-11 | ACCGTTGTAGCAATACTTCTTTGATTAGTAATAACATCACAGCTCCCAAGA | 51 |
| V8 miR-320a-3p 11-40 | GCCTAACCCGTTCAACCGATTGAGGGAGGGAAGGTAAATATTGACGGAAAT | 51 |
| V4 miR-27a-5p 40-11 | ACCGTTGTAGCAATACTTCTTTGATTAGTAATAACATCACGTCCCAAGAGC | 51 |
| V8 miR-27a-5p 11-40 | CTGACTTGTGATCAACCGATTGAGGGAGGGAAGGTAAATATTGACGGAAAT | 51 |
| V4 miR-27a-3p 40-11 | ACCGTTGTAGCAATACTTCTTTGATTAGTAATAACATCACAACCCATGGAA | 51 |
| V8 miR-27a-3p 10-40 | TTCAGTTCTCATCAACCGATTGAGGGAGGGAAGGTAAATATTGACGGAAAT | 51 |
| V4 miR-27b-5p 40-11 | ACCGTTGTAGCAATACTTCTTTGATTAGTAATAACATCACGTTCACCAATC | 51 |
| V8 miR-27b-5p 11-40 | AGCTAAGCTCTTCAACCGATTGAGGGAGGGAAGGTAAATATTGACGGAAAT | 51 |
| V4 miR-27b-3p 40-11 | ACCGTTGTAGCAATACTTCTTTGATTAGTAATAACATCACGCAGAACTTAG | 51 |
| V8 miR-27b-3p 10-40 | CCACTGTGAATCAACCGATTGAGGGAGGGAAGGTAAATATTGACGGAAAT | 50 |
| V4 miR-100-5p 40-11 | ACCGTTGTAGCAATACTTCTTTGATTAGTAATAACATCACCTGAAGAACTG | 51 |
| V8 miR-100-5p 11-40 | AATTTCAGAGGTCAACCGATTGAGGGAGGGAAGGTAAATATTGACGGAAAT | 51 |
| V4 miR-100-3p 40-11 | ACCGTTGTAGCAATACTTCTTTGATTAGTAATAACATCACAACCCCTATCAC | 52 |

| | | |
|---|---|---|
| V8 miR-100-3p 11-40 | GATTAGCATTAATCAACCGATTGAGGGAGGGAAGGTAAATATTGACGGAAAT | 52 |
| V4 miR-381-5p 40-11 | ACCGTTGTAGCAATACTTCTTTGATTAGTAATAACATCACTGTTAATGCTA | 51 |
| V8 miR-381-5p 11-40 | ATATGTAGGAGTCAACCGATTGAGGGAGGGAAGGTAAATATTGACGGAAAT | 51 |
| V4 miR-381-3p 40-11 | ACCGTTGTAGCAATACTTCTTTGATTAGTAATAACATCACAGTGAATTCTA | 51 |
| V8 miR-381-3p 11-40 | CCAGTGCCATATCAACCGATTGAGGGAGGGAAGGTAAATATTGACGGAAAT | 51 |
| V4 miR-422a 40-11 | ACCGTTGTAGCAATACTTCTTTGATTAGTAATAACATCACTCACATAGGAAT | 52 |
| V8 miR-422a 11-40 | AAAAAGCCATATCAACCGATTGAGGGAGGGAAGGTAAATATTGACGGAAAT | 51 |
| V4 miR-26a-5p 40-11 | ACCGTTGTAGCAATACTTCTTTGATTAGTAATAACATCACCGCCACGGCTC | 51 |
| V8 miR-26a-5p 11-40 | CAATCCCTATATCAACCGATTGAGGGAGGGAAGGTAAATATTGACGGAAAT | 51 |
| V4 miR-26a-1-3p 40-11 | ACCGTTGTAGCAATACTTCTTTGATTAGTAATAACATCACTTCATGGCTTC | 51 |
| V8 miR-26a-1-3p 11-40 | CATCCCTACATTCAACCGATTGAGGGAGGGAAGGTAAATATTGACGGAAAT | 51 |
| V4 miR-30e-5p 40-11 | ACCGTTGTAGCAATACTTCTTTGATTAGTAATAACATCACAACCCATGGAA | 51 |
| V8 miR-30e-5p 11-40 | TTCAGTTCTCATCAACCGATTGAGGGAGGGAAGGTAAATATTGACGGAAAT | 51 |
| V4 miR-92a-1-5p 40-12 | ACCGTTGTAGCAATACTTCTTTGATTAGTAATAACATCACACAACCAGCTA | 51 |
| V8 miR-92a-1-5p 11-40 | AGACACTGCCATCAACCGATTGAGGGAGGGAAGGTAAATATTGACGGAAAT | 51 |
| V4 miR-92a-3p 40-11 | ACCGTTGTAGCAATACTTCTTTGATTAGTAATAACATCACTCACAAGTTAG | 51 |
| V8 miR-92a-3p 11-40 | GGTCTCAGGGATCAACCGATTGAGGGAGGGAAGGTAAATATTGACGGAAAT | 51 |
| V4 miR-145-5p 40-12 | ACCGTTGTAGCAATACTTCTTTGATTAGTAATAACATCACACAACCAGCTA | 51 |
| V8 miR-145-5p 11-40 | AGACACTGCCATCAACCGATTGAGGGAGGGAAGGTAAATATTGACGGAAAT | 51 |
| V4 miR-145-3p 40-11 | ACCGTTGTAGCAATACTTCTTTGATTAGTAATAACATCACTCACAAGTTAG | 51 |
| V8 miR-145-3p 11-40 | GGTCTCAGGGATCAACCGATTGAGGGAGGGAAGGTAAATATTGACGGAAAT | 51 |
| V4 miR-200c-5p 40-11 | ACCGTTGTAGCAATACTTCTTTGATTAGTAATAACATCACAGCTCCCAAGA | 51 |
| V8 miR-200c-5p 11-40 | GCCTAACCCGTTCAACCGATTGAGGGAGGGAAGGTAAATATTGACGGAAAT | 51 |
| V4 miR-200c-3p 40-12 | ACCGTTGTAGCAATACTTCTTTGATTAGTAATAACATCACGTCCCAAGAGC | 51 |
| V8 miR-200c-3p 11-40 | CTGACTTGTGATCAACCGATTGAGGGAGGGAAGGTAAATATTGACGGAAAT | 51 |
| V4 miR-423-5p 40-12 | ACCGTTGTAGCAATACTTCTTTGATTAGTAATAACATCACAACCCATGGAA | 51 |
| V8 miR-423-5p 11-40 | TTCAGTTCTCATCAACCGATTGAGGGAGGGAAGGTAAATATTGACGGAAAT | 51 |
| V4 miR-423-3p 40-12 | ACCGTTGTAGCAATACTTCTTTGATTAGTAATAACATCACCTGAAGAACTG | 51 |
| V8 miR-423-3p 11-40 | AATTTCAGAGGTCAACCGATTGAGGGAGGGAAGGTAAATATTGACGGAAAT | 51 |
| V4 miR-29a-5p 40-11 | ACCGTTGTAGCAATACTTCTTTGATTAGTAATAACATCACAACCCCTATCAC | 52 |
| V8 miR-29a-5p 11-40 | GATTAGCATTAATCAACCGATTGAGGGAGGGAAGGTAAATATTGACGGAAAT | 52 |
| V4 miR-29a-3p 40-11 | ACCGTTGTAGCAATACTTCTTTGATTAGTAATAACATCACTGTTAATGCTA | 51 |
| V8 miR-29a-3p 11-40 | ATATGTAGGAGTCAACCGATTGAGGGAGGGAAGGTAAATATTGACGGAAAT | 51 |
| V4 miR-29b-1-5p 40-12 | ACCGTTGTAGCAATACTTCTTTGATTAGTAATAACATCACAGTGAATTCTA | 51 |
| V8 miR-29b-1-5p 12-40 | CCAGTGCCATATCAACCGATTGAGGGAGGGAAGGTAAATATTGACGGAAAT | 51 |
| V4 miR-29b-2-5p 40-11 | ACCGTTGTAGCAATACTTCTTTGATTAGTAATAACATCACTCACATAGGAAT | 52 |
| V8 miR-29b-2-5p 11-40 | AAAAAGCCATATCAACCGATTGAGGGAGGGAAGGTAAATATTGACGGAAAT | 51 |
| V4 miR-29b-3p 40-12 | ACCGTTGTAGCAATACTTCTTTGATTAGTAATAACATCACCGCCACGGCTC | 51 |
| V8 miR-29b-3p 11-40 | CAATCCCTATATCAACCGATTGAGGGAGGGAAGGTAAATATTGACGGAAAT | 51 |
| V4 miR-148a-5p 40-11 | ACCGTTGTAGCAATACTTCTTTGATTAGTAATAACATCACTTCATGGCTTC | 51 |

| | | |
|---|---|---|
| V8 miR-148a-5p 11-40 | CATCCCTACATTCAACCGATTGAGGGAGGGAAGGTAAATATTGACGGAAAT | 51 |
| V4 miR-148a-3p 40-11 | ACCGTTGTAGCAATACTTCTTTGATTAGTAATAACATCACCGCCACGGCTC | 51 |
| V8 miR-148a-3p 11-40 | CAATCCCTATATCAACCGATTGAGGGAGGGAAGGTAAATATTGACGGAAAT | 51 |
| V4 miR-455-5p 40-11 | ACCGTTGTAGCAATACTTCTTTGATTAGTAATAACATCACTTCATGGCTTC | 51 |
| V8 miR-455-5p 11-40 | CATCCCTACATTCAACCGATTGAGGGAGGGAAGGTAAATATTGACGGAAAT | 51 |
| V4 miR-455-3p 40-11 | ACCGTTGTAGCAATACTTCTTTGATTAGTAATAACATCACTTCATGGCTTC | 51 |
| V8 miR-455-3p 10-40 | CATCCCTACATTCAACCGATTGAGGGAGGGAAGGTAAATATTGACGGAAAT | 51 |
| V4 miR-181c/5p 40-11 | ACCGTTGTAGCAATACTTCTTTGATTAGTAATAACATCACACTCACCGACA | 51 |
| V8 miR-181c/5p 11-40 | GGTTGAATGTTTCAACCGATTGAGGGAGGGAAGGTAAATATTGACGGAAAT | 51 |
| V4 miR-181c/3p 40-11 | ACCGTTGTAGCAATACTTCTTTGATTAGTAATAACATCACGTCCACTCAAC | 51 |
| V8 miR-181c/3p 11-40 | GGTCGATGGTTTCAACCGATTGAGGGAGGGAAGGTAAATATTGACGGAAAT | 51 |
| V4 miR-342/5p 40-11 | ACCGTTGTAGCAATACTTCTTTGATTAGTAATAACATCACTCAATCACAGA | 51 |
| V8 miR-342/5p 10-40 | TAGCACCCCTTCAACCGATTGAGGGAGGGAAGGTAAATATTGACGGAAAT | 50 |
| V4 miR-342/3p 40-12 | ACCGTTGTAGCAATACTTCTTTGATTAGTAATAACATCACACGGGTGCGATT | 52 |
| V8 miR-342/3p 11-40 | TCTGTGTGAGATCAACCGATTGAGGGAGGGAAGGTAAATATTGACGGAAAT | 51 |

| Multiplexing detector sequences | | |
|---|---|---|
| # | Sequence | Length |
| V4 miR-15b | ACCGTTGTAGCAATACTTCTTTGATTAGTAATAACATCACTGTAAACCATG | 51 |
| V5 mir-15b 11-40 | ATGTGCTGCTAATTCGACAACTCGTATTAAATCCTTTGCCCGAACGTTATT | 51 |
| V4 miR-16 | ACCGTTGTAGCAATACTTCTTTGATTAGTAATAACATCACCGCCAATATTT | 51 |
| V6 miR-16 | ACGTGCTGCTATGGGTTATATAACTATATGTAAATGCTGATGCAAATCCAA | 51 |
| V4 miR-15a | ACCGTTGTAGCAATACTTCTTTGATTAGTAATAACATCACCACAAACCATT | 51 |
| V7 mir-15a 11-40 | ATGTGCTGCTAGTTTTAGCGAACCTCCCGACTTGCGGGAGGTTTTGAAGCC | 51 |
| V4 miR-107 | ACCGTTGTAGCAATACTTCTTTGATTAGTAATAACATCACTGATAGCCCTGT | 52 |
| V8 miR-107 | ACAATGCTGCTTCAACCGATTGAGGGAGGGAAGGTAAATATTGACGGAAAT | 51 |

| Filler sequences | | |
|---|---|---|
| # | Sequence | Length |
| V4 filler | TCTGTCCATCACGCAAATTA | 20 |
| V5 filler | AATTTTAAAAGTTTGAGTAA | 20 |
| V6 filler | TCGCAAGACAAAGAACGCGA | 20 |
| V7 filler | TCGCAAGACAAAGAACGCGA | 20 |
| V8 filler | TATTCATTAAAGGTGAATTA | 20 |

| Synthetic microRNA sequences | | |
|---|---|---|
| # | Sequence | Length |
| miR-16 | UAGCAGCACGUAAAUAUUGGCG | 22 |
| miR-107 | AGCAGCAUUGUACAGGGCUAUCA | 23 |
| miR-15a | UAGCAGCACAUAAUGGUUUGUG | 22 |
| miR-15b | UAGCAGCACAUCAUGGUUUACA | 22 |

| MicroRNA sequences for detection from total RNA | | |
|---|---|---|
| # | Sequence | Length |
| miR-16 | UAGCAGCACGUAAAUAUUGGCG | 22 |
| miR-107 | AGCAGCAUUGUACAGGGCUAUCA | 23 |
| miR-15a | UAGCAGCACAUAAUGGUUUGUG | 22 |
| miR-15b | UAGCAGCACAUCAUGGUUUACA | 22 |
| miR-34a-5p | UGGCAGUGUCUUAGCUGGUUGU | 22 |
| miR-125b-1-3p | ACGGGUUAGGCUCUUGGGAGCU | 22 |
| miR-125b-2-3p | UCACAAGUCAGGCUCUUGGGAC | 22 |
| miR-146a-5p | UGAGAACUGAAUUCCAUGGGUU | 22 |
| miR-146a-3p | CCUCUGAAAUUCAGUUCUUCAG | 22 |
| miR-155-5p | UUAAUGCUAAUCGUGAUAGGGGUU | 24 |
| miR-155-3p | CUCCUACAUAUUAGCAUUAACA | 22 |
| miR-183-5p | UAUGGCACUGGUAGAAUUCACU | 22 |
| miR-135a-5p | UAUGGCUUUUUAUUCCUAUGUGA | 23 |
| miR-135a-3p | UAUAGGGAUUGGAGCCGUGGCG | 22 |
| miR-135a-2-3p | AUGUAGGGAUGGAAGCCAUGAA | 22 |
| miR-9-5p | UCUUUGGUUAUCUAGCUGUAUGA | 23 |
| miR-9-3p | AUAAAGCUAGAUAACCGAAAGU | 22 |
| miR-128-1-5p | CGGGGCCGUAGCACUGUCUGAGA | 23 |
| miR-128-3p | UCACAGUGAACCGGUCUCUUU | 21 |
| miR-197-5p | CGGGUAGAGAGGGCAGUGGGAGG | 23 |
| miR-197-3p | UUCACCACCUUCUCCACCCAGC | 22 |
| miR-511-5p | GUGUCUUUUGCUCUGCAGUCA | 21 |
| miR-511-3p | AAUGUGUAGCAAAAGACAGA | 20 |
| miR-320a-5p | GCCUUCUCUUCCCGGUUCUUCC | 22 |
| miR-320a-3p | AAAAGCUGGGUUGAGAGGGCGA | 22 |
| miR-27a-5p | AGGGCUUAGCUGCUUGUGAGCA | 22 |
| miR-27a-3p | UUCACAGUGGCUAAGUUCCGC | 21 |
| miR-27b-5p | AGAGCUUAGCUGAUUGGUGAAC | 22 |
| miR-27b-3p | UUCACAGUGGCUAAGUUCUGC | 21 |
| miR-100-5p | AACCCGUAGAUCCGAACUUGUG | 22 |
| miR-100-3p | CAAGCUUGUAUCUAUAGGUAUG | 22 |

| | | |
|---|---|---|
| miR-381-5p | AGCGAGGUUGCCCUUUGUAUAU | 22 |
| miR-381-3p | UAUACAAGGGCAAGCUCUCUGU | 22 |
| miR-422a | ACUGGACUUAGGGUCAGAAGGC | 22 |
| miR-26a-5p | UUCAAGUAAUCCAGGAUAGGCU | 22 |
| miR-26a-1-3p | CCUAUUCUUGGUUACUUGCACG | 22 |
| miR-30e-5p | UGUAAACAUCCUUGACUGGAAG | 22 |
| miR-92a-1-5p | AGGUUGGGAUCGGUUGCAAUGCU | 23 |
| miR-92a-3p | UAUUGCACUUGUCCCGGCCUGU | 22 |
| miR-145-5p | GUCCAGUUUUCCCAGGAAUCCCU | 23 |
| miR-145-3p | GGAUUCCUGGAAAUACUGUUCU | 22 |
| miR-200c-5p | CGUCUUACCCAGCAGUGUUUGG | 22 |
| miR-200c-3p | UAAUACUGCCGGGUAAUGAUGGA | 23 |
| miR-423-5p | UGAGGGGCAGAGAGCGAGACUUU | 23 |
| miR-423-3p | AGCUCGGUCUGAGGCCCCUCAGU | 23 |
| miR-29a-5p | ACUGAUUUCUUUUGGUGUUCAG | 22 |
| miR-29a-3p | UAGCACCAUCUGAAAUCGGUUA | 22 |
| miR-29b-1-5p | GCUGGUUUCAUAUGGUGGUUUAGA | 24 |
| miR-29b-2-5p | CUGGUUUCACAUGGUGGCUUAG | 22 |
| miR-29b-3p | UAGCACCAUUUGAAAUCAGUGUU | 23 |
| miR-148a-5p | AAAGUUCUGAGACACUCCGACU | 22 |
| miR-148a-3p | UCAGUGCACUACAGAACUUUGU | 22 |
| miR-455-5p | UAUGUGCCUUUGGACUACAUCG | 22 |
| miR-455-3p | GCAGUCCAUGGGCAUAUACAC | 21 |
| miR-181c-5p | AACAUUCAACCUGUCGGUGAGU | 22 |
| miR-181c-3p | AACCAUCGACCGUUGAGUGGAC | 22 |
| miR-342-5p | AGGGGUGCUAUCUGUGAUUGA | 21 |
| miR-342-3p | UCUCACACAGAAAUCGCACCCGU | 23 |

| Other oligonucleotides | | |
|---|---|---|
| **#** | **Sequence** | **Length** |
| Blocking | ACGGTCTCATGGCCCTTCAATC | 22 |
| BtsCI cut site oligo | CTACTAATAGTAGTAGCATTAACATCCAATAAATCATACA | 40 |
